# Dog and Human Brains Represent Human Emotional Bodies with Asymmetric Encoding of Happiness and Anger

**DOI:** 10.64898/2026.10.28.762252

**Authors:** Laura V. Cuaya, Raúl Hernández-Pérez, Magdalena Boch, Janosch Rumpf, Azalea Reyes-Aguilar, Ludwig Huber, Claus Lamm

**Affiliations:** Department of Cognition, Emotion, and Methods in Psychology, Faculty of Psychology, University of Vienna, Vienna, Austria; Messerli Research Institute, Department of Interdisciplinary Life Sciences, University of Veterinary Medicine Vienna, Veterinärplatz 1, 1210, Vienna, Austria; Faculty of Psychology, National Autonomous University of Mexico, Mexico City, México

**Keywords:** bodily emotion perception, bodily happiness, bodily anger, visual social perception, representational similarity analysis, comparative neuroscience, dog fMRI

## Abstract

Dogs live in an environment rich in human emotional signals, yet most of what we know about their visual emotion perception comes from faces. Our study shifts this focus to the body, a central channel of emotional communication in dogs and a source of social information when they observe human emotional expressions. Here, we investigated how the brains of dogs and humans represent emotions conveyed by human bodies. Using fMRI, we examined 24 awake companion dogs and 40 humans viewing videos of bodily happiness, anger, and neutral actions, alongside phase-scrambled controls. Both dogs and humans showed occipito-temporal sensitivity to human bodies, with emotional modulation in the right caudal suprasylvian gyrus in dogs and the right extrastriate body area in humans. Multivariate response patterns also differentiated bodily happiness from anger in both dogs and humans. Emotion-specific RSA further showed a similar representational asymmetry: happiness showed consistent category-specific neural patterns, whereas anger showed no comparable category effect but could still be differentiated from happiness and neutral actions. Despite this shared representational asymmetry, the anatomical distribution differed, with happiness represented across more distributed regions in dogs, whereas anger differentiation involved a broader cortical network in humans. No corresponding effects were found for phase-scrambled controls. Together, these findings provide the first direct neural evidence of how the dog brain represents affective information conveyed by human bodies, while leaving open whether emotions expressed by conspecific bodies evoke similar neural representations. More broadly, our results suggest similar representational relationships between bodily emotions in dogs and humans, despite differences in the anatomical distribution of these effects, while highlighting the role of human bodily expressions in dogs’ social cognition, and advancing our understanding of the neural basis of emotional communication across species.

---

Emotion perception lies at the roots of social cognition across species. Human social environments are saturated with emotional cues, conveyed through faces, voices, postures, and movements. Living in a human-centred social world, dogs build upon attunement to human emotions to navigate social interactions, foster bonds, and cooperate with their caregivers and other persons. Using visual stimuli, it has been shown that dogs respond differentially to emotional and neutral human faces (Deputte & Doll, 2011; Hori et al., 2011; Nagasawa et al., 2011). Moreover, they discriminate happy faces from those expressing anger, fear, sadness, and disgust (Albuquerque et al., 2016; Buttelmann & Tomasello, 2013; Lazzaroni et al., 2023; Merola et al., 2014; Morisaki et al., 2009; Müller et al., 2015) and their behavior can be modulated by emotional faces (Buttelmann & Tomasello, 2013; Merola et al., 2014). Notably, Müller et al. (2015) showed that, when discriminating between happy and angry faces, dogs relied on the emotional expression of the images rather than on local facial cues. Echoing these behavioral findings, neuroimaging studies have identified brain regions in dogs that are responsive to human emotional faces (Karl, Boch, Zamansky, et al., 2020; Kujala et al., 2020; Thompkins et al., 2018, 2021). More specific insight was recently provided by a functional magnetic resonance imaging (fMRI) study revealing a distributed cluster, including the rostral sylvian gyrus, prorean gyrus, piriform cortex, and caudate nucleus, whose patterns of activity distinguish happiness from anger, sadness, and fear in human faces (Hernández-Pérez et al., 2026). Together, the behavioural and neuroimaging findings converge to establish that dogs can visually discriminate human emotions expressed in faces.

In humans, however, the perception of emotions does not rely exclusively on facial expressions but considers the whole body. Behavioural evidence shows that people can recognize emotions from bodily expressions, which can sometimes be more informative than faces (Atkinson et al., 2004; Aviezer et al., 2012; De Gelder, 2009). This sensitivity emerges early in development: infants match bodily expressions to emotional voices and show differential neural responses to emotional postures, with both effects dependent on upright presentation (Heck et al., 2018; Missana et al., 2014). Evidence from other primates also supports the relevance of bodily signals. Chimpanzees preferentially attend to agonistic interactions even when facial information is removed, but not when the videos are scrambled (Kano & Tomonaga, 2010), while macaques show stronger responses in the superior temporal sulcus to angry than to fearful or neutral conspecific body postures (De Gelder & Partan, 2009). Moreover, this neural sensitivity extends beyond conspecifics, with macaque body-selective patches responding more strongly to fearful than neutral bodies from macaques, humans, and cats (Taubert et al., 2022). Human neuroimaging identifies distributed visual, parietal, and affective regions sensitive to bodies, as well as stronger neural responses to emotionally congruent face–body displays than to isolated faces (Bonda et al., 1996; Hadjikhani & De Gelder, 2003; Peelen & Downing, 2005; Poyo Solanas et al., 2018). Within this broader network, the extrastriate body area (EBA) shows emotional modulation, supporting the idea that emotional meaning can shape processing in body-selective visual cortex (Peelen et al., 2007). Although body-selective responses often show a right-hemispheric bias (Downing et al., 2001; Smekal et al., 2026), evidence for hemispheric asymmetries in emotional and social perception remains mixed, with reports of right-EBA effects (Ross et al., 2019; Van den Stock et al., 2014), left-EBA effects (Gandolfo et al., 2024; Ren et al., 2024), and no significant hemispheric differences in emotional modulation (Peelen et al., 2007). Altogether, behavioural and neural evidence indicates that bodies convey rich affective information and that humans reliably process emotional information from bodies, providing a comparative reference for assessing whether other species, such as dogs, rely on bodily information for emotion perception.

Curiously, emerging behavioural and neural evidence suggests not only that dogs can perceive human bodily expressions of emotions, but also that, unlike humans, they may anchor their perception of human emotions more primarily in bodily expressions than in facial expressions. Reflecting this idea, an eye-tracking study showed that when dogs viewed videos of whole humans expressing emotions, they primarily attended to bodies rather than to faces, even though both were presented simultaneously, while human participants exhibited the opposite pattern (Correia-Caeiro et al., 2021). Resonating with this idea, an fMRI study presented images of headless human and dog bodies and found that the dog caudal suprasylvian gyrus (cSSG), a region located in the temporal cortex, showed preferential responses to bodies over faces and objects, and that greater neural pattern similarity for human and dog bodies than for objects was identified in the right cSSG, although emotional content was not manipulated (Boch et al., 2023). These findings point to bodily expressions as a salient social source in dogs’ visual perception of human emotions.

However, knowledge of how these processes and brain responses shape the perception of human emotion in dogs is absent. Moreover, it remains unexplored how the neural processing of human bodily emotions in dogs compares with that in humans. In the present dog-human comparative fMRI study, we therefore investigated the neural correlates of how dogs process human bodily emotions. First, we tested whether the cSSG in dogs and the EBA in humans, previously identified as body-sensitive regions, are modulated by the emotional content conveyed in bodily expressions. Second, we aimed to identify brain regions that discriminate between bodily happiness and bodily anger in dogs and humans. Third, we tested whether any brain regions show emotion-specific representations of bodily happiness and bodily anger in dogs and humans. We hypothesized that (i) the cSSG in dogs and the EBA in humans will, in each hemisphere, exhibit stronger responses to bodies than to phase-scrambled controls, and that their responses will be modulated by emotional content, with higher responses to bodily emotions (happiness and anger) than to bodily neutral; (ii) temporo-occipital regions in dogs and humans will differentiate between bodily happiness and bodily anger, both in their univariate and multivariate response patterns; and (iii) emotion-specific representational patterns will be detectable in dogs and humans for bodily happiness and bodily anger, while phase-scrambled control conditions will not show specific representational patterns will not show comparable emotion-specific representational patterns.

## Methods

### Participants

#### Dog participants

We included 24 companion dogs (Table 1) in the study (11 females), aged between 2.42 and 10.13 years (mean age = 5.84 years, SD = 2.00). The sample included a variety of breeds, specifically six Australian Shepherds, six mixed-breed, three Border Collies, two Labrador Retrievers, two Nova Scotia Duck Tolling Retrievers, one Belgian Shepherd, one Goldendoodle, one Kleiner Münsterländer, one Leisha, and one Magyar Vizsla. All dogs were trained to remain still inside an MR scanner while fully awake and unrestrained, and they were free to leave the sessions at any time (Karl, Boch, Virányi, et al., 2020). All experimental procedures were approved by the institutional ethics and animal welfare commission at the University of Veterinary Medicine Vienna (ETK-06/06/2017), in accordance with Good Scientific Practice guidelines and national legislation. The study complies with the ARRIVE guidelines (Kilkenny et al., 2010). Informed consent was obtained from the dogs’ caregivers.

**Table 1.** Dog participants.

| ID | Name | Breed | Sex | Age<sup>1</sup> |
|---|---|---|---|---|
| 01 | Edgar | Mix | Male | 5.8 |
| 02 | Moon | Australian Shepherd | Female | 3.4 |
| 03 | Minou | Goldendoodle | Female | 7.5 |
| 04 | Milo | Australian Shepherd | Male | 5.9 |
| 05 | Nikita | Australian Shepherd | Female | 6.8 |
| 06 | Jack | Australian Shepherd | Male | 3.4 |
| 07 | Mia | Mix | Female | 6.5 |
| 08 | Knopfi | Australian Shepherd | Male | 8.5 |
| 09 | Zserbo | Kleiner Münsterländer | Male | 5.3 |
| 10 | Asta | Border Collie | Female | 6.5 |
| 11 | Finlay | Border Collie | Male | 2.4 |
| 12 | Striezi | Mix | Male | 6.6 |
| 13 | Kono | Belgian Shepherd | Female | 5.1 |
| 14 | Henry | Mix | Male | 5.0 |
| 15 | Kalea | Labrador Retriever | Female | 4.7 |
| 16 | Alaska | Leisha | Female | 5.5 |
| 17 | Kelly | Australian Shepherd | Female | 3.1 |
| 18 | Black Velvet | Labrador Retriever | Female | 10.1 |
| 19 | Kazuo | Nova Scotia Duck Tolling Retriever | Male | 3.5 |
| 20 | Ori | Nova Scotia Duck Tolling Retriever | Female | 7.3 |
| 21 | Ace | Border Collie | Male | 6.3 |
| 22 | Mailo | Mix | Male | 10.1 |
| 23 | Timo | Mix | Male | 6.5 |
| 24 | Danny | Magyar Viszla | Male | 4.5 |

<sup>1</sup> Age in years at the beginning of their participation in the experiment

#### Human participants

We included 40 healthy adults (24 women), aged between 18-43 years (mean age = 25.9 years, SD = 6.14). Participants were recruited through the Vienna CogSciHub: Study Participant Platform (SPP), which uses the hroot software (Bock et al., 2014). All participants provided informed consent based on the GDPR template (OBC-GDPR-ULT 1.1.0) recommended by the Open Brain Consent working group (Bannier et al., 2021) and received €30 compensation for their participation. All human experimental procedures were approved by the Ethics Committee of the University of Vienna (Reference Number: 01389) and conducted in accordance with the Declaration of Helsinki.

### Data acquisition

#### Dog participants

MRI data were obtained using a custom-made 16-channel (k9) head coil specifically designed for the anatomy of the dog head (Guran et al., 2023), used in a 3 T Siemens Skyra MR system (Siemens Medical, Erlangen, Germany). We acquired whole-brain blood-oxygen-level-dependent (BOLD) images using a 2-fold multiband (MB) accelerated gradient-echo echo-planar imaging (EPI) sequence for the functional scans with the following parameters: voxel size = 1.5 x 1.5 x 2 mm3, 20% gap; repetition time (TR) = 1000 ms; echo time (TE) = 38 ms; flip angle = 61°; field of view (FOV) = 144 x 144 x 58 mm3; 24 axial slices (interleaved acquisition, descending order) with a posterior-to-anterior phase-encoding direction; 280 volumes. In a separate scan session prior to functional data collection, we acquired a T1-weighted structural image with the following parameters: isotropic voxel size = 0.7 mm3; TR = 2100 ms; TE = 3.3 ms; FOV = 230 x 230 x 165 mm3. Participants wore noise-protecting earplugs. During acquisitions, the dog trainer remained inside the scanner room to monitor and handle the dog out of their sight. After each functional run, the dogs were given a rest period outside the scanner. Functional images were successful when they met the following criteria: 1) translational movement within a maximum of 3 mm in any direction; 2) rotational movement within a maximum of 3° in any direction; and 3) a maximum of 33% of volumes with framewise displacement (FD) > 0.5 mm (Power et al., 2012). Each participant completed at least two runs (range = 2 to 6, median = 4). In total, we acquired 88 successful runs (Table S1).

#### Human participants

MRI data were obtained using a 32-channel head coil in a 3 T Siemens Skyra MR system (Siemens Medical, Erlangen, Germany). BOLD images were acquired using a 4-fold MB accelerated gradient-echo EPI sequence for the functional scans with the following parameters: voxel size = 2 x 2 x 2 mm3, 20% gap; TR = 1200 ms; TE = 34 ms; flip angle = 66°; FOV = 192 x 192 x 124.8 mm3; 52 axial slices (interleaved acquisition, ascending order) with a posterior-to-anterior phase-encoding direction; 234 volumes. A T1-weighted structural image was acquired at the end of the scanning session with the following parameters: isotropic voxel size = 0.94 mm3; TR = 2300 ms; TE = 2.29 ms; FOV = 240 x 240 x 165.4 mm3. All four functional runs were acquired within a single scanning session and were alternated with runs from a separate experiment. In total, we acquired 160 runs.

### Stimuli

To increase the ecological validity of the stimuli, we: 1) presented videos instead of images, as the expression of emotion has an important dynamic component; 2) presented the videos in color; 3) provided actors with a common situation in which the target emotion towards a dog could arise naturally (e.g., for anger: “*You just finished tidying up your home, and as you turn around, the dog has tracked muddy paw prints all over the freshly cleaned floor and furniture.*”; and for happiness: “*The dog performs a new trick perfectly after you’ve been training it for weeks, surprising you with their accomplishment.*”) and instructed them to perform as naturally as possible, focusing on their body expression; and 4) positioned the camera 80 cm from the floor to approximate the dogs’ point of view (Karl et al., 2021).

We recorded twelve actors (six actresses) in a single session while they performed three types of conditions: happiness, anger, and the action of changing a light bulb with a neutral emotion. Each actor received compensation between 30 and 50 euros, depending on the time required for filming. Actors were aware of the aim of the study and the post-processing that would be performed on their recorded videos; they signed an informed consent. All actors were unfamiliar to the dogs who participated in this study. The videos (resolution of 1280 x 720 pixels and 30 frames/s) were recorded using a GoPro Hero 9 camera under standardized conditions, in the same room to maintain similar visual properties across recordings, a one-way ANOVA confirmed that there were no significant differences (all p > 0.05) in brightness, hue, saturation, or contrast across the three conditions. Professional photo lamps were used to prevent visual distractions caused by shadows during the videos. We followed guidelines of previous studies on creating stimuli for emotional human bodily expressions (Atkinson et al., 2004; De Gelder & Van den Stock, 2011). Actors were instructed to portray the three conditions with naturalistic body movements. They had the freedom to interpret the emotions naturally, providing minimal guidance on the context in which these emotions could typically be experienced toward a dog. Each take lasted at least 8 s, and we recorded at least two takes, varying depending on the actor’s and experimenter’s judgment during the filming. The recorded videos were edited to adjust the duration to 4 s, focusing on the peak of the emotion. We selected two 4 s videos per condition per actor (i.e., 36 videos). Then, facial information was selectively removed from each video by manually applying a blur effect to the head region in the first frame and distributing it across all frames using automated facial tracking in Adobe After Effects (Adobe Inc., San Jose, CA). For the action videos, we selected the take in which all the flow of the action was clear. For the emotional videos (happiness and anger), we conducted an online survey in which adult humans (n = 20, 12 women, mean age = 32.7 years, range 20 – 45 years old) were asked to rate the valence and the intensity of each of the 24 emotional videos. Responses for both questions were given on a continuous scale. The valence of all videos was correctly identified by the participants (i.e., happiness videos labeled as positive valence and anger videos labeled as negative valence), and we selected the videos with the highest identification scores.

To control for low-level visual features, we created two additional video categories by applying phase-scrambling to the happiness and anger videos. Phase-scrambled videos have been used as control conditions in fMRI studies across species, including marmosets (Zanini et al., 2023), humans (Rajimehr et al., 2024), and dogs (Boch, Karl, et al., 2024), as they preserve low-level visual properties while disrupting high-level visual content. Phase-scrambled videos were generated in Python by computing the two-dimensional discrete Fourier transform of each frame of the videos and applying a random translation to the phase of each frequency. Each phase-scrambled frame was then reassembled in the same order as the original video. The translation kept constant across the three-color channel (RGB) and across frames, ensuring that luminance, color, and motion energy matched exactly those of the original videos, while the visual content in each frame appeared scrambled. All videos were resized to 800 × 1080 pixels and positioned at the center of the MR screen to appear in the participant’s visual field.

### Design

We used an event-related design (Figure 1). Each event consisted of a 4 s soundless video. In each run, we presented six events from each of the five categories: two bodily emotion categories, happiness and anger; and three control categories: bodily neutral (i.e., an action performed with neutral emotionality, to control for the presence of a moving human body without emotional content) and phase-scrambled happiness and anger videos (to control for low-level visual properties). Events were presented in a semi-random order (with a maximum of two consecutive events of the same category) using PsychoPy (Peirce, 2007). Each event was separated by a baseline averaging 5 s (ranging 3.5 s to 7 s in increments of 0.5 s), during which dogs observed a gray screen with a white cross. Each run lasted 280 s and ended with a 10 s baseline. We created four different runs with variation of the sequence of actors and emotions shown. Each included videos of the same six actors (three actresses, three actors) across all the conditions. The order of presentations for the runs was counterbalanced across participants.

**Figure 1.**
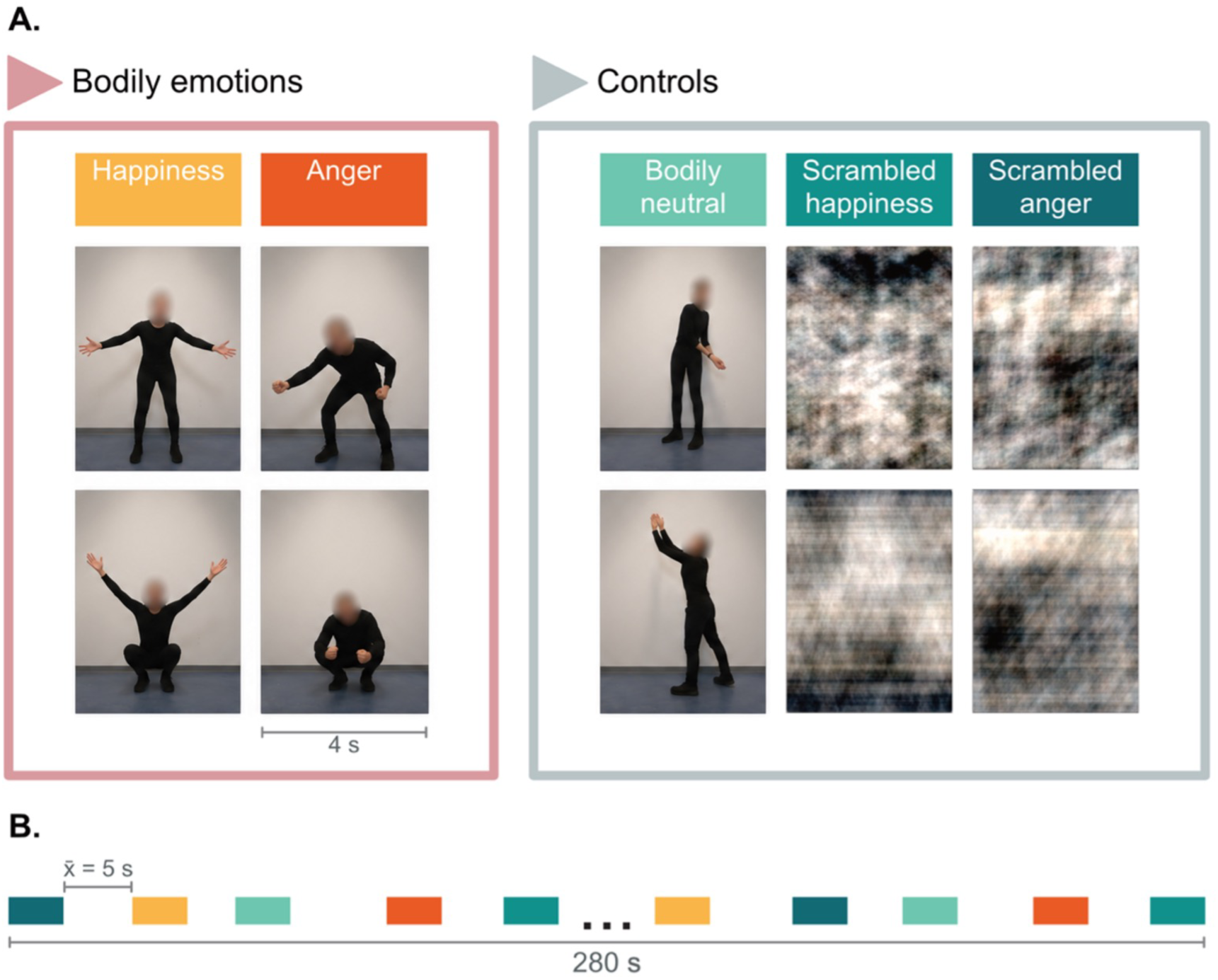
Experimental design. We presented five categories of 4 s videos: two bodily emotions (happiness and anger), and three control categories (bodily neutral, phase-scrambled happiness, and phase-scrambled anger). **A.** To comply with bioRxiv’s policy on identifiable human images, the example images of bodily happiness, anger, and neutral actions are computer-generated illustrations replacing frames from the original videos used as experimental stimuli. The top and bottom rows represent the five experimental conditions corresponding to stimuli originally recorded from one actress and one actor, respectively. **B.** Timeline of a run using an event-related design. Each run included six pseudo-randomized presentations of each video category, separated by a baseline interval averaging 5 s (range: 3 to 7 s). Each run featured six different actors (three of them actresses) across all categories. Each run lasted 280 s.

### Data analysis

Raw functional images were pre-processed using FSL 6.0.5 (Jenkinson et al., 2012) for both species. For dog participants, we additionally used the Dog Brain Toolkit, a custom toolkit for preprocessing and analyzing dog fMRI data (Hernández-Pérez, 2025). For dogs, we first applied motion correction and slice timing correction. We reoriented all the functional images to match the dog template (Nitzsche et al., 2019). We then calculated a mean functional image by aligning each run to the first volume of the first run, then averaged all volumes for each dog; each run was then aligned to the mean functional image using FLIRT (Jenkinson et al., 2002; Jenkinson & Smith, 2001). We then skull-stripped all runs using a binary mask manually drawn over the mean functional image. The mean functional image was transformed to the dog template (Nitzsche et al., 2019) using FLIRT, thus creating a mean normalized image (in the dog template space) for each dog. We calculated a transformation matrix from the mean image to the mean normalized image using FLIRT and applied it to each previously aligned run. The images were then smoothed using a Gaussian kernel (3 mm FWHM). For human participants, standard FSL preprocessing was applied, with functional images registered to the individual T1-weighted image and normalized to standard MNI space. The images were then smoothed using a Gaussian kernel (4 mm FWHM).

To describe the univariate brain response, we conducted a whole-brain General Linear Model (GLM) analysis using FEAT v6.00. Statistical parametric maps were generated using the linear combination of functions derived by convolving the corresponding hemodynamic response function (HRF) with the time series of each of the five conditions (bodily happiness, bodily anger, bodily neutral, phase-scrambled happiness, and phase-scrambled anger). For dogs, we used the dog-tailored HRF (Boch et al., 2021), whereas for humans, we used the standard double-gamma HRF. The first-level individual models included the six standard motion parameters as regressors of no interest; we censored the volumes with FD > 0.5 mm (dogs: 1259 volumes, corresponding to 5.1% of the total, Table S1; humans: 665 volumes, corresponding to 1.8% of the total). Individual contrast images with an implicit baseline were computed for each of the five conditions, from those contrasts we extracted the BOLD signal change using FEATQuery. We also included the following contrasts: All conditions > implicit baseline; Bodily conditions (happiness, anger, neutral) > phase-scrambled controls (happiness, anger); Bodily emotions (happiness, anger) > controls (bodily neutral, phase-scrambled happiness, phase-scrambled anger); Bodily happiness > bodily anger, bodily neutral, phase-scrambled happiness; Bodily anger > bodily happiness, bodily neutral, phase-scrambled anger. The resulting group statistical parametric maps were thresholded (z > 3.1 and pcluster < 0.05) using random field theory (Worsley, 2001).

To test our hypothesis that the emotional content of bodies modulates the response within established body-sensitive regions, we conducted region of interest analyses (ROI) in the bilateral caudal suprasylvian gyri (cSSG) in dogs and bilateral extrastriate body area (EBA) in humans. The cSSG was defined using a multi-scale functional dog brain atlas generated via spatially constrained spectral clustering (Hernández-Pérez et al., 2025), by selecting the functional parcels encompassing the coordinates corresponding to the left and right cSSG (20,-26, 4 and-20,-24, 2), previously identified as a body-sensitive region in dogs (Boch et al., 2023), from the segmentation solution containing 140 parcels (N=140). The EBA was defined using a multi-scale functional human brain atlas generated via spatially constrained spectral clustering (Craddock et al., 2012). We selected the functional parcels encompassing the peak coordinates of EBA (±48.9,-69.8, 0.5) based on the average coordinates reported across seventeen studies (Shultz et al., 2011), from the segmentation solution containing 200 parcels (N=200). The 200-parcel solution was chosen because it provides an optimal compromise between functional specificity and anatomical interpretability (Craddock et al., 2012). We performed a 2 (Hemisphere: left, right) × 5 (Condition: bodily happiness, bodily anger, bodily neutral, phase-scrambled happiness, phase-scrambled anger) repeated-measures ANOVA on the average percent BOLD signal change extracted from the individual contrast images for each condition.

To describe the multivariate brain response, we employed Representational Similarity Analysis (RSA). First, we obtained subject-specific activation patterns (beta estimates) for each of the five conditions (bodily happiness, bodily anger, bodily neutral, phase-scrambled happiness, phase-scrambled anger) from the GLM. These beta estimates reflected the response to each condition relative to the implicit baseline. To assess the relationship between the response patterns, we calculated a neural dissimilarity matrix (DSM) per participant. Pairwise dissimilarities were computed using the crossnobis (cross-validated Mahalanobis) distance estimator. This metric is a reliable, unbiased estimate of dissimilarity as it leverages independent measurements (e.g., separate experimental runs) for cross-validation, which effectively removes noise components that are common to the measurements (Schütt et al., 2023). We implemented RSA in the bodily emotions functional mask resulting from the GLM whole-brain contrast Bodily emotions (happiness, anger) > controls (bodily neutral, phase-scrambled happiness, phase-scrambled anger), using a searchlight approach (Kriegeskorte et al., 2006). For each voxel in the brain, a spherical searchlight (radius = 3 voxels in dogs; 4 voxels in humans) were centered on that voxel. Using all voxels contained within the sphere, we computed a 5 x 5 neural DSM by calculating the pairwise crossnobis distances for all condition pairs. To identify brain regions where the neural similarity response pattern matched a theoretical model, we created correlation maps. In these maps, the value assigned to each center voxel was the Kendall’s rank correlation coefficient τA between the locally computed neural DSM (from the searchlight analysis) and a predefined theoretical model DSM. This process resulted in one similarity map per participant. The similarity maps were averaged across all participants to create a group-level model similarity map. To account for the spatial dependence inherent in the searchlight approach and to minimize assumptions about statistical inference, we employed a random permutation method (Stelzer et al., 2013). Specifically, the labels for the five conditions were randomly shuffled to generate a random neural DSM for each participant. This random DSM was then correlated with the predefined theoretical model DSM, using Kendall’s rank correlation coefficient τA to produce random similarity maps. We then averaged these maps to create a random group-level similarity map. This procedure was repeated 10,000 times. The resulting random maps were averaged to estimate the mean correlation and standard deviation expected by chance. These parameters were then used to convert the correlation values at each center voxel into z-scores. The resulting z-maps were thresholded to retain only voxels with z > 3.1. The same threshold was applied to the random group-level similarity map. Based on these thresholded random maps, we derived the cluster size distribution expected by chance and used it to exclude clusters smaller than would be expected under the null distribution (p < 0.05).

We constructed ten theoretical models to test specific hypotheses regarding the bodily emotional information represented in dogs and humans. Five models focused on bodily emotion conditions (models 1, 3, 5, 7, and 9), while the other five were corresponding phase-scrambled control models used to test whether the representational relationships observed for emotional bodily videos could be explained by low-level visual features. The first six models shared a within-condition similarity requirement: videos within the same condition of interest were expected to elicit similar neural response patterns.

1) Emotion Valence model. It assumed high within-condition similarity for bodily happiness and for bodily anger, and high dissimilarity between bodily happiness and bodily anger.
2) Phase-scrambled Emotion Valence model. The same representational structure was tested using phase-scrambled happiness and phase-scrambled anger.

The subsequent emotion-specific models and their phase-scrambled controls, in addition to requiring high within-condition similarity, required high dissimilarity between the target condition and the two other conditions, to identify brain regions in which response patterns represented the target condition across stimuli within condition and differed from those elicited by the other two conditions.

3) Bodily Happiness model. It assumed high within-condition similarity for bodily happiness and high dissimilarity between bodily happiness and both bodily anger and bodily neutral.
4) Phase-scrambled Happiness model. The same representational structure was tested after replacing bodily happiness and bodily anger with their phase-scrambled counterparts.
5) Bodily Anger model. It assumed high within-condition similarity for bodily anger and high dissimilarity between bodily anger and both bodily happiness and bodily neutral.
6) Phase-scrambled Anger model. The same representational structure was tested after replacing bodily anger and bodily happiness with their phase-scrambled counterparts.

The final four models tested a different representational structure and did not require high within-condition similarity. Instead, they tested whether the target condition was represented as distinct from both other conditions, while the two non-target conditions were represented as similar to each other. Thus, these models aimed to identify brain regions in which the target condition occupied a distinct position in representational space, even if stimuli within that condition did not elicit highly similar response patterns.

7) Bodily Happiness Distinctiveness model. It assumed high dissimilarity between bodily happiness and both bodily anger and bodily neutral, and low dissimilarity between bodily anger and bodily neutral.
8) Phase-scrambled Happiness Distinctiveness model. The same representational structure was tested after replacing bodily happiness and bodily anger with their phase-scrambled counterparts.
9) Bodily Anger Distinctiveness model. It assumed high dissimilarity between bodily anger and both bodily happiness and bodily neutral, and low dissimilarity between bodily happiness and bodily neutral.
10) Phase-scrambled Anger Distinctiveness model. The same representational structure was tested after replacing bodily anger and bodily happiness with their phase-scrambled counterparts.

Finally, we assessed cross-species correspondence in stimulus-pair representational dissimilarities between dog and human ROIs identified by the corresponding RSA models. For each ROI, pairwise representational dissimilarities were extracted from correlation-distance searchlight maps (1 − Pearson’s r, radius = 3 voxels for dogs and radius = 4 voxels for humans). The dissimilarity value at each ROI’s searchlight centre was extracted for every stimulus pair specified by the corresponding RSA model, generating a dissimilarity vector for each participant, session, and run. For each dog and human ROI comparison, Pearson correlations were calculated between the corresponding stimulus-pair dissimilarities, separately for bodily happiness and anger pairs.

## Results

Table 2 presents the results of the five whole-brain GLM contrasts. The contrast All conditions > implicit baseline showed three clusters in dogs, primarily located in temporo-occipital regions. The largest cluster had its peak z-value in the right mid suprasylvian gyrus. In humans, the same contrast showed nine clusters distributed across occipital, parietal, temporal, and frontal regions. The largest cluster had its peak in the right occipital pole.

**Table 2.** GLM contrasts.

| Contrast and brain regions | Coordinates<sup>1</sup> |  |  | z-value | Cluster size |
|---|---|---|---|---|---|
|  | x | y | z |  |  |
| **Dog participants** |  |  |  |  |  |
| **All conditions > implicit baseline** |  |  |  |  |  |
| R mid suprasylvian gyrus | 15 | -26 | 13 | 7.81 | 2156 |
| L mesencephalon | -5 | -18 | 1 | 4.96 | 27 |
| R diencephalon | 5 | -16 | 1 | 4.65 | 19 |
| **Bodily conditions (happiness, anger, neutral) > phase-scrambled controls (happiness, anger)** |  |  |  |  |  |
| R caudal sylvian gyrus | 19 | -14 | 5 | 6.05 | 470 |
| L caudal suprasylvian gyrus | -23 | -26 | 1 | 5.83 | 444 |
| **Bodily emotions (happiness, anger) > controls (bodily neutral, phase-scrambled happiness, phase-scrambled anger)** |  |  |  |  |  |
| R rostral sylvian gyrus | 17 | -10 | 7 | 6.24 | 500 |
| L caudal suprasylvian gyrus | -23 | -24 | 1 | 6.54 | 465 |
| **Bodily happiness > bodily anger, bodily neutral, phase-scrambled happiness** |  |  |  |  |  |
| L rostral sylvian gyrus | -15 | -10 | 7 | 5.46 | 363 |
| R rostral sylvian gyrus | 17 | -12 | 9 | 5.3 | 277 |
| L marginal gyrus | -1 | -32 | 19 | 4.13 | 38 |
| L ectomarginal gyrus | -9 | -30 | 17 | 4.14 | 15 |
| **Bodily anger > bodily happiness, bodily neutral, phase-scrambled anger** |  |  |  |  |  |
| R mid suprasylvian gyrus | 15 | -26 | 15 | 5.13 | 145 |
| R rostral ectosylvian gyrus | 17 | -10 | 9 | 5.74 | 96 |
| L caudal sylvian gyrus | -21 | -12 | 7 | 4.84 | 60 |
| L caudal suprasylvian gyrus | -17 | -26 | 11 | 5.15 | 59 |
| L caudal suprasylvian gyrus | -21 | -24 | 1 | 4.44 | 27 |
| L marginal gyrus | -1 | -34 | 19 | 4.6 | 17 |
| **Human participants** |  |  |  |  |  |
| **All conditions > implicit baseline** |  |  |  |  |  |
| R occipital pole / primary visual cortex | 20 | -94 | -2 | 8.31 | 26474 |
| R precentral gyrus | 44 | 0 | 46 | 6.05 | 1267 |
| L superior parietal lobule | -24 | -58 | 54 | 5.23 | 765 |
| R parahippocampal gyrus | 18 | -30 | -6 | 6.68 | 350 |
| L parahippocampal gyrus | -18 | -30 | -4 | 7.48 | 281 |
| L amygdala / anterior parahippocampal region | -22 | -2 | -26 | 5.09 | 209 |
| L inferior frontal gyrus | -48 | 28 | -6 | 4.57 | 190 |
| L precentral gyrus | -34 | -4 | 50 | 5.16 | 160 |
| L middle temporal gyrus | -46 | -30 | 20 | 4.23 | 65 |
| **Bodily conditions (happiness, anger, neutral) > phase-scrambled controls (happiness, anger)** |  |  |  |  |  |
| R lateral occipital cortex | 46 | -78 | -8 | 7.29 | 4280 |
| L lateral occipital cortex | -46 | -78 | -2 | 6.5 | 2665 |
| R occipital pole | 14 | -94 | -4 | 4.72 | 382 |
| L parahippocampal gyrus | -18 | -4 | -20 | 4.45 | 284 |
| L inferior occipital cortex | -18 | -88 | -16 | 5.26 | 272 |
| R precuneus | 4 | -58 | 28 | 4.18 | 238 |
| R inferior frontal gyrus | 56 | 26 | 6 | 5.13 | 229 |
| R temporal pole / anterior temporal cortex | 28 | 0 | -22 | 4.65 | 196 |
| L inferior frontal gyrus | -56 | 18 | -2 | 4.86 | 189 |
| L occipital fusiform gyrus | -44 | -58 | -22 | 4.45 | 147 |
| R cerebellum | 30 | 8 | -46 | 4.05 | 82 |
| L middle temporal gyrus | -54 | 4 | -22 | 4.39 | 76 |
| L posterior temporal / MTG – STG border | -58 | -16 | -10 | 4.62 | 63 |
| **Bodily emotions (happiness, anger) > controls (bodily neutral, phase-scrambled happiness, phase-scrambled anger)** |  |  |  |  |  |
| R lateral occipital cortex (inferior division) | 48 | -76 | -8 | 7.88 | 11579 |
| L lateral occipital cortex / occipital fusiform | -46 | -78 | -16 | 7.43 | 6197 |
| L parahippocampal gyrus | -18 | -4 | -22 | 7.07 | 2235 |
| R precuneus / posterior cingulate cortex | 2 | -56 | 24 | 6.03 | 1003 |
| R superior parietal lobule | 30 | -40 | 60 | 5.34 | 648 |
| L superior parietal lobule | -28 | -44 | 66 | 5.17 | 386 |
| L parahippocampal gyrus | -14 | -32 | -4 | 5.46 | 179 |
| R medial frontal cortex | 6 | 56 | 24 | 4.47 | 171 |
| R precuneus dorsal / medial superior | 10 | 2 | 72 | 4.81 | 165 |
| L precentral gyrus | -42 | 2 | 58 | 4.58 | 160 |
| R precentral gyrus | 44 | 2 | 50 | 4.36 | 139 |
| L cerebellar crus I | -18 | -70 | -36 | 4.69 | 136 |
| L medial temporal / posterior parahippocampal region | -4 | -6 | -8 | 4.35 | 96 |
| **Bodily happiness > bodily anger, bodily neutral, phase-scrambled happiness** |  |  |  |  |  |
| R lateral occipital cortex | 52 | -64 | 2 | 7.75 | 7239 |
| L lateral occipital cortex | -48 | -80 | -2 | 7.83 | 5860 |
| R superior parietal lobule | 24 | -46 | 66 | 5.73 | 1245 |
| L superior parietal lobule | -30 | -42 | 62 | 5.65 | 1242 |
| R precuneus | 10 | 2 | 70 | 5.42 | 373 |
| R temporal pole | 30 | -2 | -26 | 4.55 | 365 |
| R inferior frontal gyrus (pars orbitalis) | 52 | 8 | -26 | 4.8 | 272 |
| R inferior frontal gyrus (pars triangularis) | 52 | 28 | 2 | 4.1 | 224 |
| R precentral gyrus | 42 | 0 | 56 | 4.49 | 217 |
| L parahippocampal gyrus | -32 | 18 | -36 | 4.37 | 151 |
| L inferior frontal gyrus (par opercularis) | -40 | 26 | -6 | 4.31 | 144 |
| L precentral gyrus | -40 | -4 | 60 | 4.47 | 127 |
| R middle frontal gyrus | 48 | 14 | 28 | 4.34 | 109 |
| L parahippocampal gyrus | -18 | -4 | -22 | 4.59 | 85 |
| **Bodily anger > bodily happiness, bodily neutral, phase-scrambled anger** |  |  |  |  |  |
| R lateral occipital cortex | 46 | -78 | -8 | 8.55 | 8124 |
| L lateral occipital cortex | -46 | -80 | 2 | 7.03 | 5135 |
| L parahippocampal gyrus | -20 | -6 | -18 | 5.31 | 1201 |
| R precuneus / posterior cingulate | 4 | -52 | 24 | 5.46 | 1045 |
| L inferior frontal gyrus (par triangularis) | -50 | 22 | -10 | 5.4 | 233 |
| R inferior Frontal gyrus | 52 | 24 | 2 | 4.56 | 220 |
| R parahippocampal gyrus | 16 | -30 | -2 | 4.89 | 153 |
| R middle frontal gyrus | 46 | 20 | 24 | 4.24 | 143 |
| L cerebellar crus I | -14 | -72 | -34 | 4.42 | 103 |
| R medial frontal cortex | 6 | 56 | 20 | 4.19 | 83 |
| L parahippocampal gyrus | -16 | -34 | -8 | 4.51 | 82 |
| L medial frontal cortex | -6 | 50 | 30 | 4.33 | 63 |

<sup>1</sup> Coordinates are given in mm according to the Nitzsche atlas for the dog data and to the stereotactic Montreal Neurological Institute (MNI) space for human data.

### Body-sensitive regions and their emotional modulation

Next, we examined the brain response to human bodies (Figure 2A). The whole-brain GLM contrast Bodily conditions (happiness, anger, neutral) > phase-scrambled controls (happiness, anger) showed two clusters in dogs, primarily located in temporo-occipital regions. The largest cluster had its peak z-value in the right caudal sylvian gyrus. Brain regions with a response to human bodies included bilateral mid and caudal suprasylvian gyri, bilateral ectosylvian gyri, bilateral sylvian gyri, bilateral caudal composite gyri, bilateral piriform lobe, left insular cortex, right straight gyrus, and right occipital gyrus. In humans, the same contrast showed 13 clusters, primarily located in occipito-temporal regions. The largest cluster had its peak z-value in the right lateral occipital cortex. Brain regions with a higher response to human bodies included bilateral occipital cortex, bilateral inferior frontal gyri, left inferior occipital cortex, left occipital fusiform gyrus, left parahippocampal gyrus, bilateral temporal regions, right precuneus, right occipital pole, and right cerebellum.

**Figure 2.**
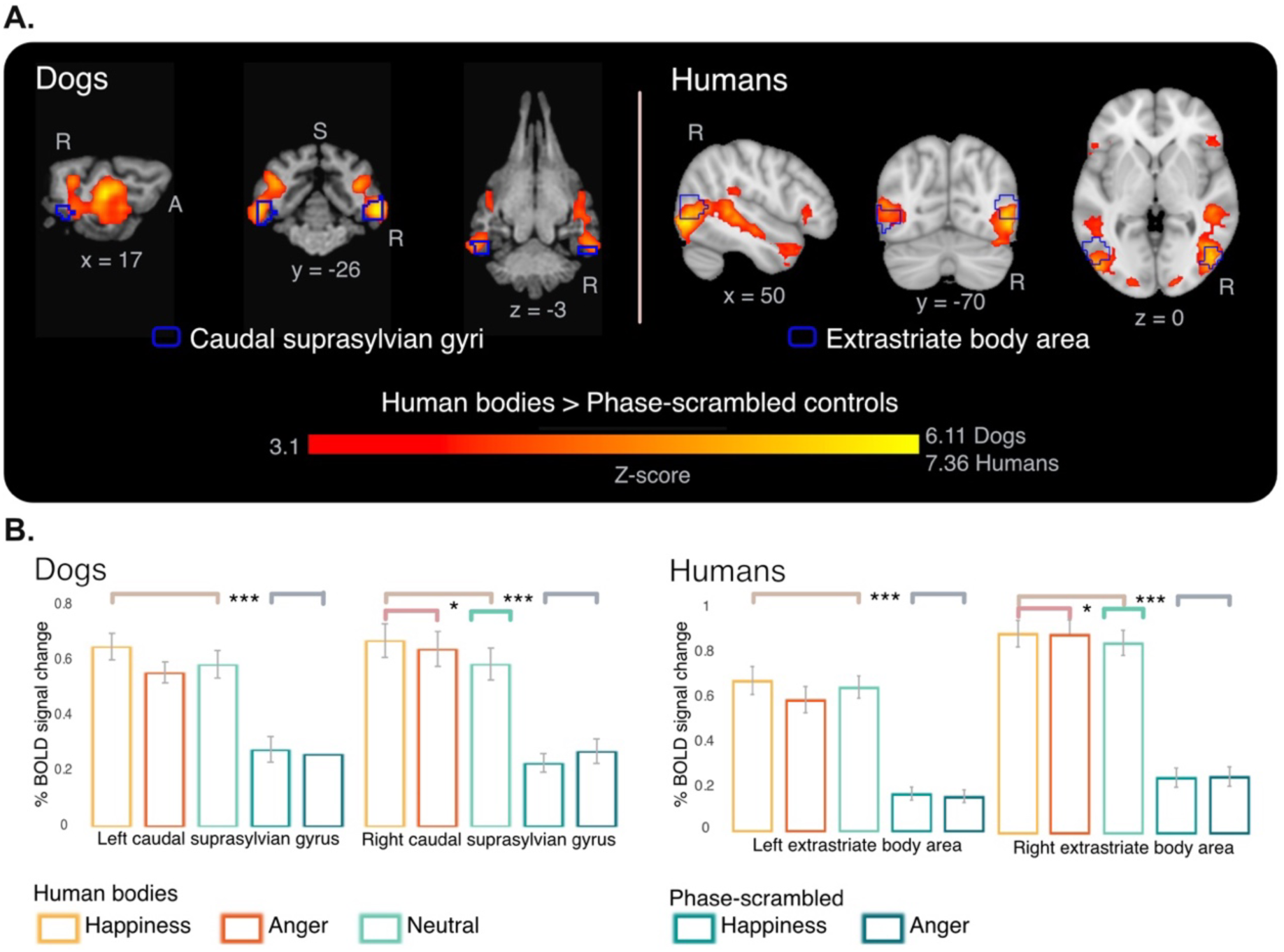
Human body-sensitive brain regions and responses in the dog caudal suprasylvian gyri (n = 24) and human extrastriate body area (n = 40). A. Whole-brain contrast Bodily conditions (happiness, anger, neutral) > phase-scrambled controls (happiness, anger) for dogs and humans, overlaid on species-specific template brains. In both species, significant responses were predominantly located in bilateral occipito-temporal regions. Outlines of the caudal suprasylvian gyri mask in dogs (Hernández-Pérez et al., 2025) and the extrastriate body area mask in humans (Craddock et al., 2012) are shown in blue. Results are thresholded at z > 3.1, pcluster < 0.05. The color bar indicates the z-score. R = right; A = anterior; S = superior. B. ROI analysis within the masks shown in panel A. Bars show mean percent BOLD signal change in each hemisphere for the five conditions. In both species and hemispheres, bodily conditions (happiness, anger, neutral) elicited higher responses than phase-scrambled controls. In the right hemisphere, bodily emotions (happiness and anger) also elicited higher responses than bodily neutral in dogs and humans. Vertical lines represent standard error. *** = p < 0.001, * = p < 0.05.

To assess emotional modulation within the body-sensitive region of interest of each species, we examined BOLD responses across five conditions (Figure 2B, Table 3). In dogs, the analysis in the bilateral caudal suprasylvian gyri (cSSG) revealed no significant main effect of Hemisphere, F(1, 23) = 0.12, p = 0.737, partial η² = 0.00; a significant main effect of Condition, F(2.46, 56.47) = 52.75, p < 0.001, partial η² = 0.70 (Greenhouse–Geisser-corrected); and no significant Hemisphere × Condition interaction, F(3.27, 75.24) = 2.19, p = 0.091, partial η² = 0.09. Planned contrasts showed that, in both the left and right cSSG, bodily conditions (happiness, anger, neutral) elicited higher responses than phase-scrambled controls (happiness, anger), both pFDR < 0.001 (FDR-corrected within each hemisphere). The contrast Bodily emotions (happiness, anger) vs. bodily neutral was significant in the right cSSG (pFDR = 0.039), but not in the left cSSG (pFDR = 0.568), indicating an emotional modulation in the right hemisphere on top of the bilateral body sensitivity in cSSG. In humans, the corresponding analysis in the bilateral extrastriate body area (EBA) revealed a significant main effect of Hemisphere, F(1, 39) = 10.09, p = 0.003, partial η² = 0.21, indicating overall stronger responses in the right hemisphere; a significant main effect of Condition, F(1.55, 60.50) = 182.57, p < 0.001, partial η² = 0.82 (Greenhouse–Geisser-corrected); and a significant Hemisphere × Condition interaction, F(1.21, 47.09) = 8.57, p = 0.003, partial η² = 0.18. Planned contrasts showed that, in both the left and right EBA, bodily conditions (happiness, anger, neutral) elicited higher responses than phase-scrambled controls (happiness, anger), both pFDR < 0.001 (FDR-corrected within each hemisphere). The contrast Bodily emotions (happiness, anger) vs. bodily neutral was significant in the right EBA (pFDR = 0.014), but not in the left EBA (pFDR = 0.528), indicating an emotional modulation in the right hemisphere on top of the bilateral body sensitivity in EBA.

**Table 3.** Planned contrasts in the dog caudal suprasylvian gyri (cSSG) and the human extrastriate body area (EBA)

| Hemisphere | Estimate | SE | df | 95% CI lower | 95% CI upper | t | p |
|---|---|---|---|---|---|---|---|
| **Dog participants** |  |  |  |  |  |  |  |
| **Bodily conditions vs. Phase-scrambled controls** |  |  |  |  |  |  |  |
| Left | 0.327 | 0.039 | 23 | 0.246 | 0.409 | 8.321 | < .001 |
| **Bodily emotions vs. Bodily neutral** |  |  |  |  |  |  |  |
| Left | 0.017 | 0.030 | 23 | –0.044 | 0.079 | 0.579 | 0.568 |
| **Bodily conditions vs. Phase-scrambled controls** |  |  |  |  |  |  |  |
| Right | 0.383 | 0.041 | 23 | 0.299 | 0.467 | 9.455 | < .001 |
| **Bodily emotions vs. Bodily neutral** |  |  |  |  |  |  |  |
| Right | 0.070 | 0.032 | 23 | 0.004 | 0.137 | 2.186 | 0.039 |
| **Human participants** |  |  |  |  |  |  |  |
| **Bodily conditions vs. Phase-scrambled controls** |  |  |  |  |  |  |  |
| Left | 0.472 | 0.046 | 39 | 0.379 | 0.566 | 10.236 | < .001 |
| **Bodily emotions vs. Bodily neutral** |  |  |  |  |  |  |  |
| Left | -0.013 | 0.021 | 39 | –0.056 | 0.029 | -0.637 | 0.528 |
| **Bodily conditions vs. Phase-scrambled controls** |  |  |  |  |  |  |  |
| Right | 0.624 | 0.044 | 39 | 0.536 | 0.713 | 14.326 | < .001 |
| **Bodily emotions vs. Bodily neutral** |  |  |  |  |  |  |  |
| Right | 0.039 | 0.015 | 39 | 0.008 | 0.070 | 2.560 | 0.014 |

### Distributed whole-brain responses to bodily emotions

In dogs, the contrast Bodily emotions (happiness, anger) > controls (bodily neutral, phase-scrambled happiness, phase-scrambled anger) showed two clusters, mainly located in the temporal lobe but also involving parietal, occipital, and insular cortices. The largest cluster had its peak z-value in the right rostral sylvian gyrus. Brain regions with a higher response to bodily emotions included bilateral mid suprasylvian gyri, bilateral ectosylvian gyri, bilateral sylvian gyri, bilateral caudal composite gyri, and bilateral insular cortex. These regions defined the dog bodily emotions functional mask used in subsequent analyses. In humans, the same contrast showed thirteen clusters, predominantly located in occipito-temporal regions, with additional involvement of parietal and frontal cortices. The largest cluster had its peak z-value in the right lateral occipital cortex. Brain regions with a higher response to bodily emotions included bilateral lateral occipital cortex extending into occipital fusiform areas, medial temporal regions including the parahippocampal gyrus, bilateral superior parietal lobules, the precuneus/posterior cingulate cortex, medial frontal cortex, precentral gyri, and cerebellar regions. These results defined the human bodily emotions functional mask used in subsequent analyses.

### The univariate brain response to bodily happiness and anger

We then compared whole-brain responses to bodily happiness and bodily anger (Figure 3). For bodily happiness, in dogs, the contrast Bodily happiness > bodily anger, bodily neutral, phase-scrambled happiness showed four clusters, primarily along the temporal lobe, with the maximum peak z-value within the two biggest clusters in the bilateral rostral sylvian gyri. Brain regions with a higher response to bodily happiness included the bilateral mid and caudal suprasylvian gyri, right rostral ectosylvian gyrus, bilateral mid and caudal ectosylvian gyrus, bilateral sylvian gyri, bilateral caudal composite gyri, right marginal gyrus, bilateral ectomarginal gyri, and right splenial gyrus. In humans, the same contrast showed fourteen clusters, primarily located in occipital-parietal regions, with the maximum peak z-value within the two largest clusters in the bilateral lateral occipital cortex. Brain regions with a higher response to bodily happiness included bilateral lateral occipital cortex, bilateral superior parietal lobules, precuneus, right temporal pole, bilateral inferior frontal regions, bilateral precentral gyri, middle frontal cortex, and medial temporal regions. For bodily anger, in dogs, the contrast Bodily anger > bodily happiness, bodily neutral, phase-scrambled anger showed six clusters, also primarily along the temporal lobe, with the maximum peak z-value within the largest cluster in the right mid suprasylvian gyrus. Brain regions with a higher response to bodily anger included the bilateral mid and caudal suprasylvian gyri, right rostral and mid ectosylvian gyri, bilateral sylvian gyri, and left marginal gyrus. In humans, the same contrast showed twelve clusters, primarily located in occipito-temporal regions, with the maximum peak z-value within the two largest clusters in the bilateral lateral occipital cortex. Brain regions with a higher response to bodily anger included bilateral lateral occipital cortex, medial temporal regions including the parahippocampal gyrus, precuneus/posterior cingulate cortex, bilateral inferior frontal regions, middle and medial frontal cortices, and cerebellar regions.

**Figure 3.**
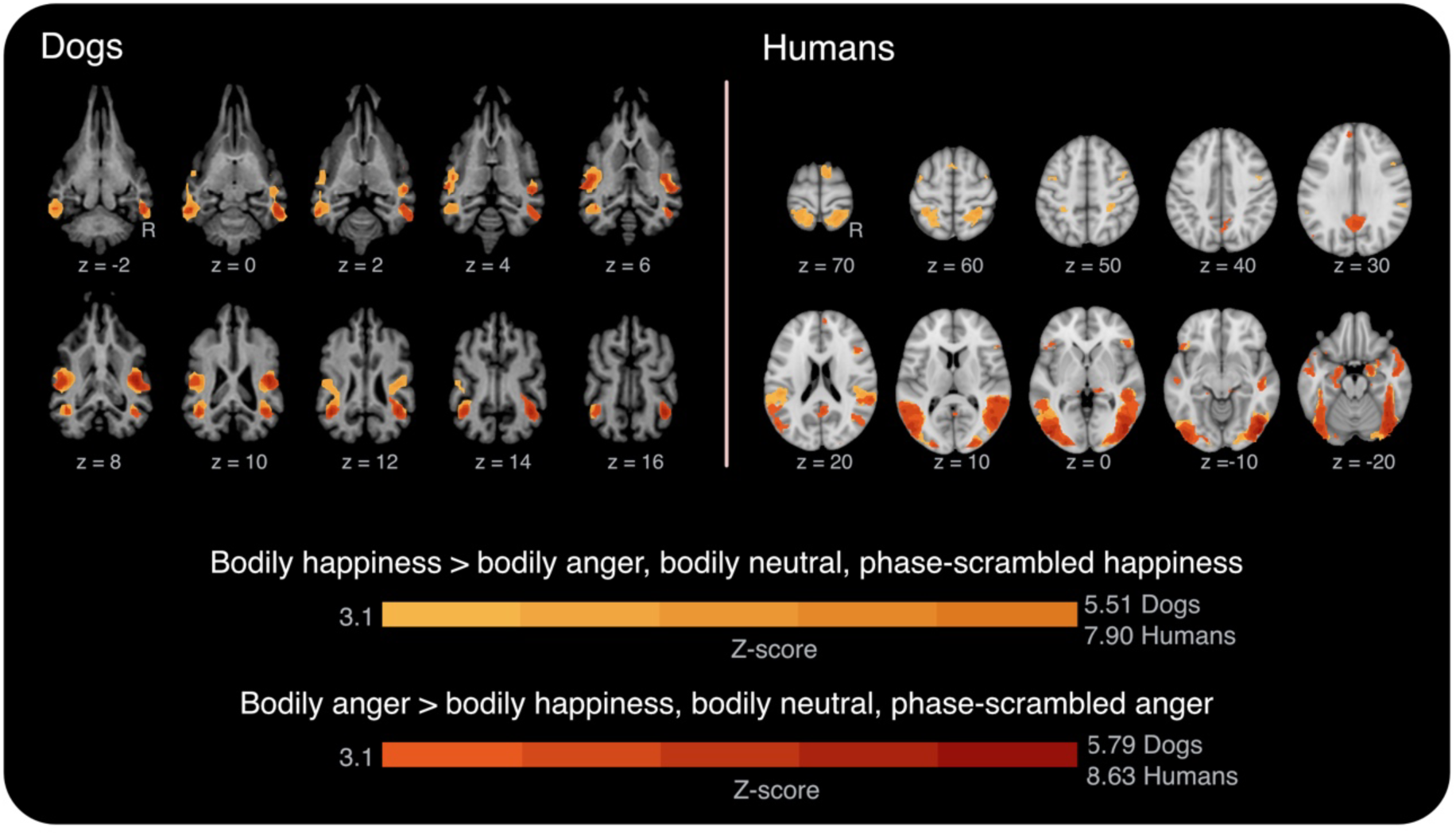
Univariate brain responses to bodily happiness and anger in dogs (n = 24) and humans (n = 40). Whole-brain contrasts (z > 3.1, pcluster < 0.05) for Bodily happiness > bodily anger, bodily neutral, phase-scrambled happiness and Bodily anger > bodily happiness, bodily neutral, phase-scrambled anger are shown across consecutive axial slices for dogs and humans. Bodily anger responses are overlaid on bodily happiness responses. R = right.

In dogs, although some differences in spatial extent and distribution were observed, such as a broader and more anterior brain response related to bodily happiness and a more posterior response to bodily anger, both contrasts showed substantial overlap. This overlap was also evident when examining the BOLD signal in 3 mm spheres centered on each of the peaks of the ten clusters (four for happiness, six for anger), where the univariate BOLD responses were generally similar for bodily happiness and anger (Figures S1 and S2). In humans, the two contrasts also showed substantial overlap, particularly within bilateral occipito-temporal cortex and visual body-sensitive regions, although this overlap was less extensive than in dogs. Differences in spatial distribution were also observed. Bodily happiness was associated with a broader dorsal and sensorimotor pattern, including bilateral superior parietal lobules, bilateral opercular regions, right superior frontal gyrus, and bilateral precentral gyri. Bodily anger showed additional responses in medial and anterior regions, including bilateral precuneus and posterior cingulate cortex, bilateral temporal poles, as well as left-lateralized frontal and temporal areas encompassing the inferior frontal gyrus, middle temporal gyrus, orbital frontal cortex, and superior frontal gyrus.

### Multivariate brain representation of bodily emotions

As a next approach, we employed multivariate analyses to assess sensitivity to emotional valence in each species. We used the Emotion Valence model to test valence-specific representations independently of bodily neutral responses (Figure 4, Table 4). We conducted a searchlight analysis within the species-specific bodily emotions functional masks, defined from the GLM whole-brain contrast Bodily emotions (happiness, anger) > controls (bodily neutral, phase-scrambled happiness, phase-scrambled anger), to identify regions where activity patterns discriminated between bodily happiness and bodily anger. This model therefore tested whether patterns of activity within brain regions selectively responsive to bodily emotional information also encoded valence-specific distinctions between positive and negative bodily expressions. In dogs, the analysis showed two clusters, with the maximum peak z-value within the largest cluster located in the right caudal ectosylvian gyrus. Brain regions where the response patterns elicited by bodily happiness differed significantly from those elicited by bodily anger included, in the right hemisphere, the rostral and caudal ectosylvian gyri and the rostral and caudal sylvian gyrus, and in the left hemisphere, the mid and caudal suprasylvian gyri. In humans, the same analysis showed one cluster, with the maximum peak z-value located in the right occipital pole. Brain regions where the response patterns elicited by bodily happiness differed significantly from those elicited by bodily anger included the right occipital pole and right lateral occipital cortex. Even though this analysis was conducted within the bodily emotions functional masks, where, by definition, the response to the phase-scrambled controls was lower, we tested the Phase-scrambled Emotion Valence model to determine whether the discrimination between emotional valences in the previous model (Emotion Valence model) could be driven by low-level visual properties in either species. We repeated the searchlight analysis to find regions where phase-scrambled happiness and phase-scrambled anger elicited distinct response patterns. No significant results were found in dogs nor humans.

**Figure 4.**
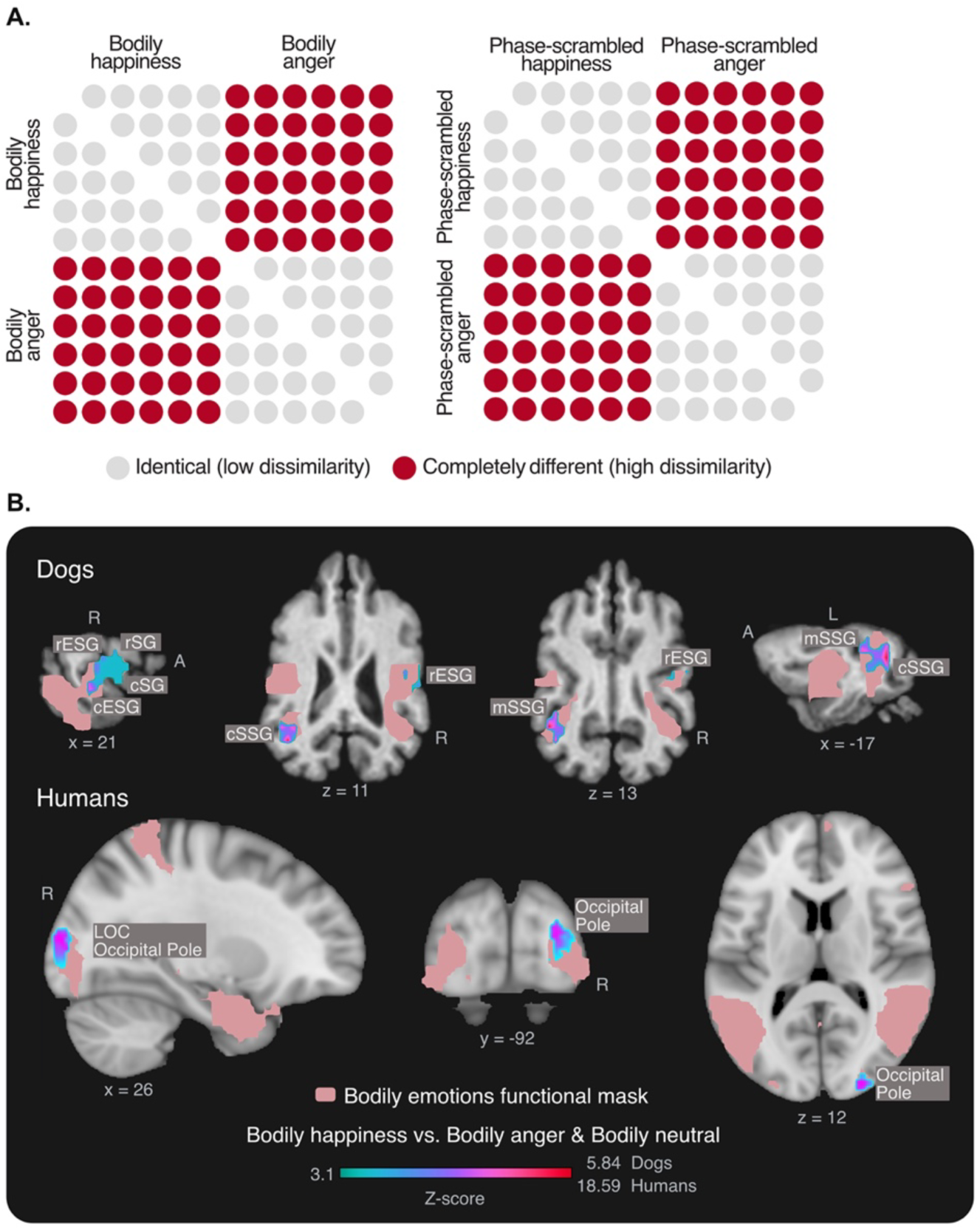
**Representational similarity analysis (RSA) results for the Emotion Valence model in dogs (n = 24) and humans (n = 40)**. **A.** Dissimilarity matrices (DSMs) representing the Emotion Valence model (left) and the Phase-scrambled Emotion Valence model (right). Matrix rows and columns correspond to the conditions (bodily happiness and bodily anger in the left panel; phase-scrambled happiness and phase-scrambled anger in the right panel). The colour code indicates the hypothesized representational distance: light grey circles denote low dissimilarity (conditions expected to produce similar patterns within the same emotion), while red circles denote high dissimilarity (conditions expected to produce distinct neural patterns between positive and negative valence). The Emotion Valence model tests for brain regions that discriminate between bodily happiness and bodily anger, while the Phase-scrambled Emotion Valence model tests for brain regions that discriminate between phase-scrambled happiness and phase-scrambled anger. **B.** RSA results for dogs and humans are overlaid on species-specific template brains. Lateral and axial views show the regions where the local neural DSM significantly correlated with the theoretical Emotion Valence DSM within the species-specific bodily emotions functional masks. Results are thresholded at z > 3.1, pcluster < 0.05. The colour bar indicates the z-score of the Kendall’s τA correlation. rESG = rostral ectosylvian gyrus; cESG = caudal ectosylvian gyrus; cSG = caudal sylvian gyrus; rSG = rostral sylvian gyrus; cSSG = caudal suprasylvian gyrus; mSSG = mid suprasylvian gyrus; R = right; A = anterior; L = left. No significant results were found for the Phase-scrambled Emotion Valence model.

**Table 4.** Regions discriminating bodily happiness and anger1.

| Cluster and brain regions | Coordinates<sup>2</sup> |  |  | z-value | Cluster size |
|---|---|---|---|---|---|
|  | x | y | z |  |  |
| **Dog participants** |  |  |  |  |  |
| **Cluster 1** |  |  |  |  | 60 |
| R caudal ectosylvian gyrus | 21 | -16 | 1 | 4.91 |  |
| R rostral ectosylvian gyrus | 17 | -8 | 11 | 4.86 |  |
| R rostral sylvian gyrus | 25 | -12 | 7 | 4.77 |  |
| **Cluster 2** |  |  |  |  | 40 |
| L mid suprasylvian gyrus | -19 | -24 | 13 | 5.77 |  |
| **Human participants** |  |  |  |  |  |
| **Cluster 1** |  |  |  |  | 152 |
| R occipital pole | 26 | -92 | 12 | 18.40 |  |
| R lateral occipital cortex, inferior division | 30 | -84 | 8 | 5.17 |  |

<sup>1</sup> Emotion Valence model, searchlight within bodily emotions functional mask.
<sup>2</sup> Coordinates are given in mm according to the Nitzsche atlas for the dog data and to the stereotactic Montreal Neurological Institute (MNI) space for human data.

After identifying the regions where patterns of response distinguished between bodily happiness and bodily anger, our next aim was to identify regions within the bodily emotions functional mask that were specifically involved in processing each emotion (Figure 5, Table 5). For the Bodily Happiness model, we searched for brain regions where blocks of the bodily happiness condition showed high within-condition similarity and high dissimilarity relative to both the bodily anger and bodily neutral conditions. In dogs, we found three clusters, with the maximum peak z-value within the biggest cluster located in the right caudal sylvian gyrus. Brain regions where the response patterns for bodily happiness were both consistent and distinct from those elicited by bodily anger or bodily neutral conditions included, in the first cluster (right hemisphere), the caudal sylvian gyrus, rostral sylvian gyrus, rostral ectosylvian gyrus, and the insular cortex; in the second cluster (left hemisphere), the mid suprasylvian gyrus and caudal suprasylvian gyrus; and in the third cluster, the right mid suprasylvian gyrus. In humans, the same model revealed one cluster, with the maximum peak z-value located in the right occipital pole. The response pattern for bodily happiness was both consistent and distinct from those elicited by bodily anger and bodily neutral conditions in the right occipital pole. To determine whether low-level visual features explained the previous results, we applied the searchlight RSA within the bodily emotions functional masks to the phase-scrambled conditions. For the Phase-scrambled Happiness model, we tested for regions where the phase-scrambled conditions elicited a distinct response pattern, characterized by high within-condition similarity and high dissimilarity relative to phase-scrambled anger and bodily neutral conditions. No significant results were found in dogs or humans. Similarly, to identify regions specifically involved in the processing of anger, we applied the same searchlight RSA approach within the bodily emotions functional masks. For the Bodily Anger model, we searched for brain regions where blocks of the bodily anger condition showed high within-condition similarity and high dissimilarity relative to both the bodily happiness and bodily neutral conditions. No significant results were found in dogs or humans, meaning that no region within the bodily emotions functional masks showed a response pattern for bodily anger that was both consistent and distinct from those elicited by bodily happiness and bodily neutral conditions. Likewise, when applying the same searchlight RSA to the phase-scrambled conditions, no significant results were found for the Phase-scrambled Anger model in dogs or humans.

**Figure 5.**
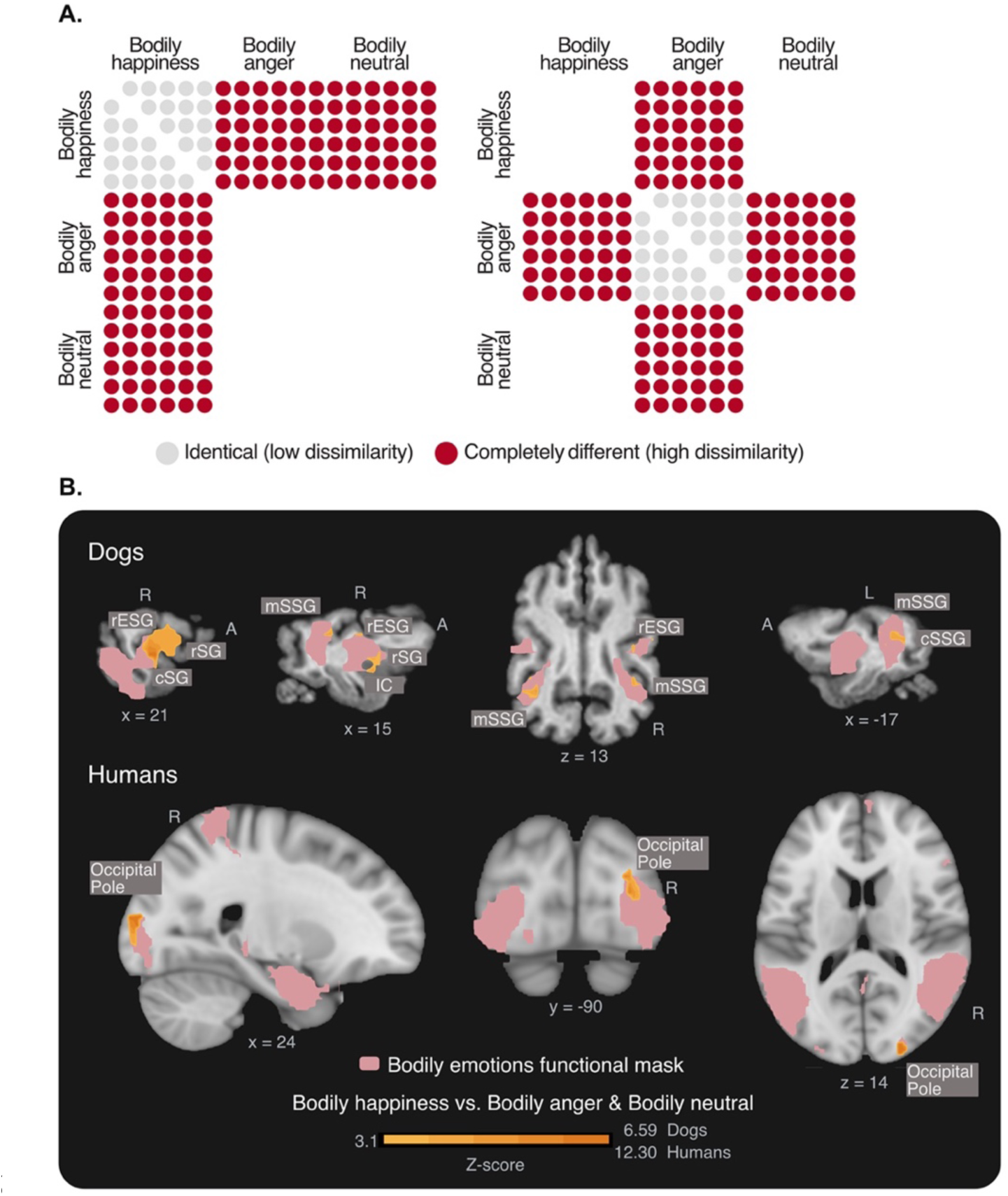
Representational similarity analysis (RSA) results for the Bodily Happiness model in dogs (n = 24) and humans (n = 40). **A.** Dissimilarity matrices (DSM) representing the Bodily Happiness model (left) and the Bodily Anger model (right). Matrix rows and columns correspond to the conditions (bodily happiness, bodily anger, and bodily neutral). The colour code indicates the hypothesized representational distance: light grey circles denote low dissimilarity (conditions expected to produce similar response patterns), while red circles denote high dissimilarity (conditions expected to produce distinct response patterns). Pairs without circles were not included in either model. The Bodily Happiness model tests for brain regions where the bodily happiness condition elicits a distinct response pattern relative to bodily anger and bodily neutral; the Bodily Anger model tests for regions where the bodily anger condition elicits a distinct response pattern relative to bodily happiness and bodily neutral. **B.** RSA results for dogs and humans are overlaid on species-specific template brains. Selected views show the regions where the local neural DSM significantly correlated with the theoretical Bodily Happiness DSM within the species-specific bodily emotions functional masks. Results are thresholded at z > 3.1, pcluster < 0.05. The colour bar indicates the z-score of the Kendall’s τA correlation. rESG = rostral ectosylvian gyrus; cSG = caudal sylvian gyrus; rSG = rostral sylvian gyrus; mSSG = mid suprasylvian gyrus; IC = insular cortex; cSSG = caudal suprasylvian gyrus; R = right; A = anterior; L = left. No significant results were found for the Bodily Anger model in either species.

**Table 5.** Regions responsive to bodily happiness1.

| Cluster and brain regions | Coordinates<sup>2</sup> |  |  | z-value | Cluster<br>size |
|---|---|---|---|---|---|
|  | x | y | z |  |  |
| **Dog participants** |  |  |  |  |  |
| **Cluster 1** |  |  |  |  | 71 |
| R caudal sylvian gyrus | 21 | -12 | 3 | 6.53 |  |
| R insular cortex | 15 | -6 | 1 | 4.79 |  |
| R rostral ectosylvian gyrus | 21 | -10 | 11 | 4.38 |  |
| **Cluster 2** |  |  |  |  | 18 |
| L caudal suprasylvian gyrus | -17 | -28 | 11 | 5.23 |  |
| L mid suprasylvian gyrus | -17 | -20 | 13 | 4.10 |  |
| **Cluster 3** |  |  |  |  | 6 |
| R mid suprasylvian gyrus | 15 | -20 | 13 | 4.46 |  |
| **Human participants** |  |  |  |  |  |
| **Cluster 1** |  |  |  |  | 121 |
| R occipital pole | 24 | -90 | 14 | 12.18 |  |

<sup>1</sup> Bodily Happiness model, searchlight within bodily emotions functional mask.
<sup>2</sup> Coordinates are given in mm according to the Nitzsche atlas for the dog data and to the stereotactic Montreal Neurological Institute (MNI) space for human data.

Then, we conducted a whole-brain searchlight RSA using the Distinctiveness models (Figure 6, Table 6). Whereas the Bodily Happiness and Bodily Anger models tested whether the target emotion elicited a response pattern characterized by high within-condition similarity and high dissimilarity relative to each of the other conditions, the Distinctiveness models focused on between-condition representational relationships. Specifically, they tested whether bodily anger or bodily happiness was represented as distinct from the other two conditions, while the other two conditions were expected to be more similar to each other, without requiring high within-condition similarity for the target emotion. For the Bodily Anger Distinctiveness model, in dogs, the analysis revealed one significant cluster, with the maximum peak z-value located in the right rostral composite gyrus. In humans, the same model revealed two significant clusters, with the maximum peak z-value within the largest cluster located in the left lateral occipital cortex. Brain regions showing the Bodily Anger Distinctiveness pattern included bilateral lateral occipital cortex and additional occipital, parietal, sensorimotor, and insular regions within the largest cluster, while the second cluster included the right paracingulate gyrus, right superior frontal gyrus, and right frontal pole. No significant results were found in dogs or humans for the Bodily Happiness Distinctiveness model, the Phase-scrambled Happiness Distinctiveness model, or the Phase-scrambled Anger Distinctiveness model. This suggests that, in dogs and humans, the neural representation of bodily anger as distinct from happiness and neutral was detectable only when high within-condition similarity was not required, whereas no comparable effect was found for bodily happiness or the phase-scrambled controls.

**Figure 6.**
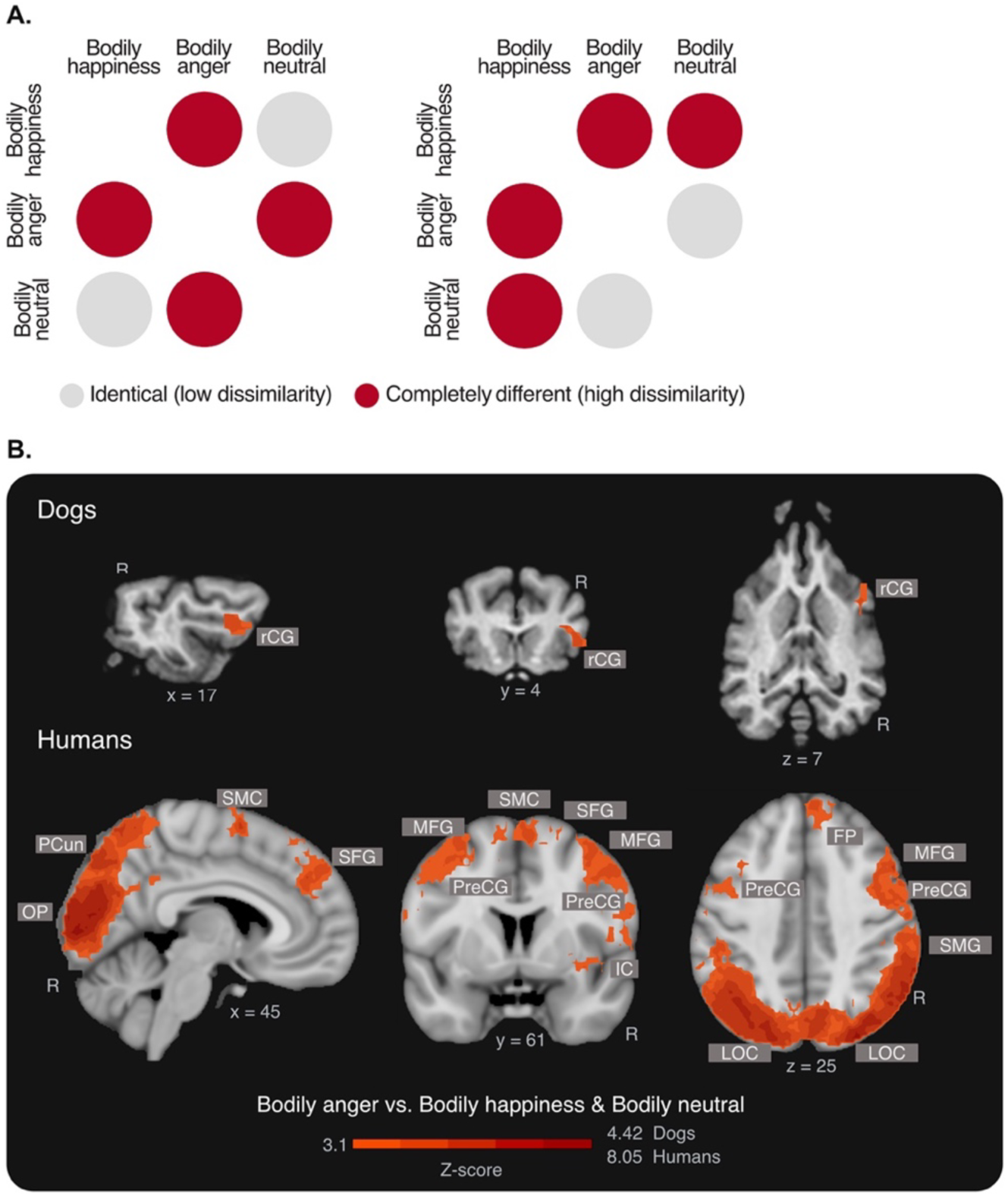
Representational similarity analysis (RSA) results for the Bodily Anger Distinctiveness model in dogs (n = 24) and humans (n = 40). A. Dissimilarity matrices (DSMs) representing the theoretical Bodily Anger Distinctiveness model (left) and Bodily Happiness Distinctiveness model (right). Matrix rows and columns correspond to the experimental conditions (bodily happiness, bodily anger, bodily neutral). Red circles denote high dissimilarity, and light grey circles denote low dissimilarity. In the Bodily Anger Distinctiveness model, bodily anger is expected to elicit a response pattern distinct from both bodily happiness and bodily neutral, whereas bodily happiness and bodily neutral are expected to be more similar. In the Bodily Happiness Distinctiveness model, bodily happiness is expected to elicit a response pattern distinct from both bodily anger and bodily neutral, whereas bodily anger and bodily neutral are expected to be more similar. B. Whole-brain RSA results for dogs and humans, overlaid on species-specific template brains. In dogs, the Bodily Anger Distinctiveness model revealed a significant cluster in the right rostral composite gyrus, where the local neural DSM significantly correlated with the theoretical model. In humans, the same model revealed distributed significant effects encompassing occipital, parietal, sensorimotor, insular, and frontal regions. Results are thresholded at z > 3.1, pcluster < 0.05. The colour bar indicates the z-score of the Kendall’s τA correlation. No significant results were found for the Bodily Happiness Distinctiveness model. rCG = rostral composite gyrus; FP = frontal pole; IC = insular cortex; LOC = lateral occipital cortex; MFG = middle frontal gyrus; OP = occipital pole; PCun = precuneous; PreCG = precentral gyrus; SFG = superior frontal gyrus; SMC = supplementary motor cortex; SMG = supramarginal gyrus; R = right.

**Table 6.** Regions responsive to bodily anger1.

| Cluster and brain regions | Coordinates<sup>2</sup> |  |  | z-value | Cluster size |
|---|---|---|---|---|---|
|  | x | y | z |  |  |
| **Dog participants** |  |  |  |  |  |
| **Cluster 1** |  |  |  |  | 38 |
| R rostral composite gyrus | 13 | 6 | 9 | 4.37 |  |
| R rostral composite gyrus | 21 | 4 | 3 | 3.66 |  |
| **Human participants** |  |  |  |  |  |
| **Cluster 1** |  |  |  |  | 35993 |
| L lateral occipital cortex | -38 | -81 | 26 | 7.91 |  |
| L lateral occipital cortex | -24 | -81 | 22 | 7.94 |  |
| R lateral occipital cortex | 60 | -61 | 19 | 7.91 |  |
| **Cluster 2** |  |  |  |  | 750 |
| R paracingulate gyrus | 6 | 43 | 34 | 5.46 |  |
| R superior frontal gyrus | 4 | 52 | 39 | 5.08 |  |
| R frontal pole | 14 | 51 | 35 | 5.02 |  |

<sup>1</sup> Bodily Anger Distinctiveness model, whole-brain searchlight.
<sup>2</sup> Coordinates are given in mm according to the Nitzsche atlas for the dog data and to the stereotactic Montreal Neurological Institute (MNI) space for human data.

Finally, to directly assess cross-species correspondence in the representation of bodily emotions, we compared stimulus-pair dissimilarities between dog and human ROIs identified by the corresponding RSA models Table 7, Figure S3). Cross-species correlations were higher for pairs of bodily happiness stimuli than for pairs of bodily anger stimuli in 20 of the 26 ROI comparisons (76.9%), with median Pearson correlation coefficients of 0.510 and 0.369, respectively. However, this pattern was observed in only four of the eight comparisons involving ROIs identified by the Emotion Valence model.

**Table 7.**
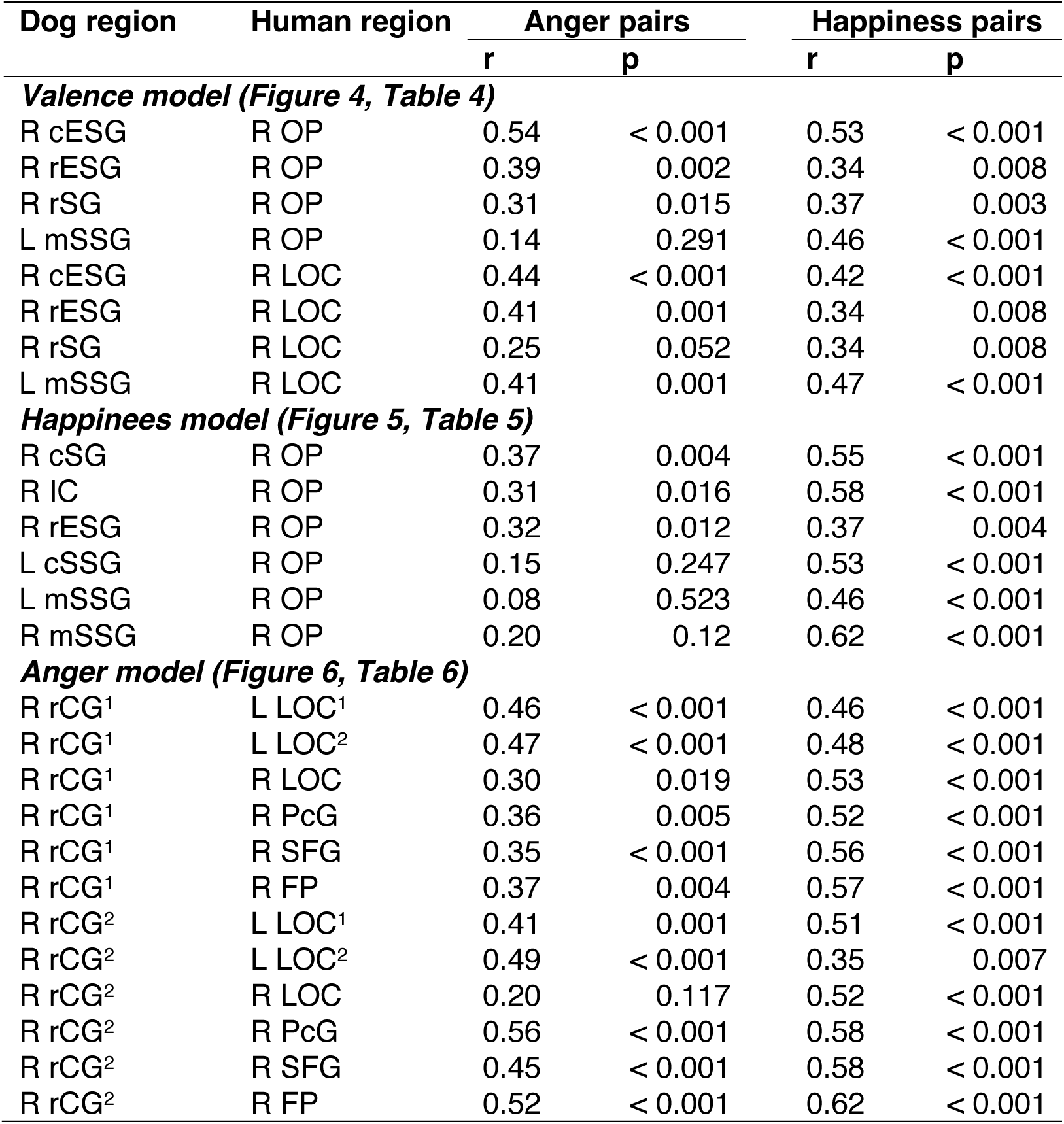
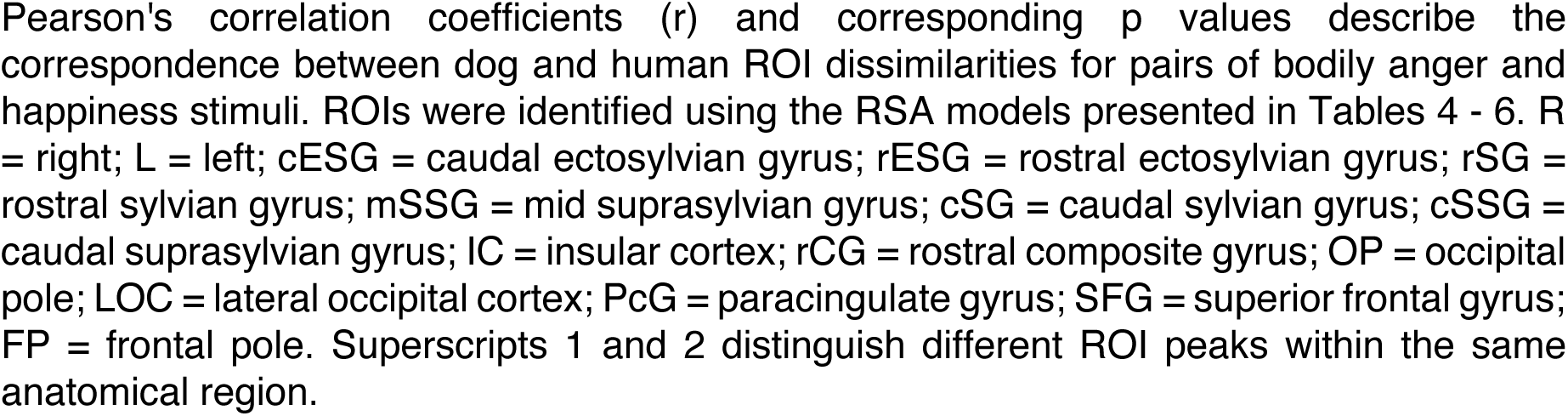
Cross-species correlations of stimulus-pair dissimilarities in dog and human brain regions.

| Dog region | Human region | Anger pairs |  | Happiness pairs |  |
|---|---|---|---|---|---|
|  |  | r | p | r | p |
| ***Valence model (Figure 4, Table 4)*** |  |  |  |  |  |
| R cESG | R OP | 0.54 | < 0.001 | 0.53 | < 0.001 |
| R rESG | R OP | 0.39 | 0.002 | 0.34 | 0.008 |
| R rSG | R OP | 0.31 | 0.015 | 0.37 | 0.003 |
| L mSSG | R OP | 0.14 | 0.291 | 0.46 | < 0.001 |
| R cESG | R LOC | 0.44 | < 0.001 | 0.42 | < 0.001 |
| R rESG | R LOC | 0.41 | 0.001 | 0.34 | 0.008 |
| R rSG | R LOC | 0.25 | 0.052 | 0.34 | 0.008 |
| L mSSG | R LOC | 0.41 | 0.001 | 0.47 | < 0.001 |
| ***Happinees model (Figure 5, Table 5)*** |  |  |  |  |  |
| R cSG | R OP | 0.37 | 0.004 | 0.55 | < 0.001 |
| R IC | R OP | 0.31 | 0.016 | 0.58 | < 0.001 |
| R rESG | R OP | 0.32 | 0.012 | 0.37 | 0.004 |
| L cSSG | R OP | 0.15 | 0.247 | 0.53 | < 0.001 |
| L mSSG | R OP | 0.08 | 0.523 | 0.46 | < 0.001 |
| R mSSG | R OP | 0.20 | 0.12 | 0.62 | < 0.001 |
| ***Anger model (Figure 6, Table 6)*** |  |  |  |  |  |
| R rCG<sup>1</sup> | L LOC<sup>1</sup> | 0.46 | < 0.001 | 0.46 | < 0.001 |
| R rCG<sup>1</sup> | L LOC<sup>2</sup> | 0.47 | < 0.001 | 0.48 | < 0.001 |
| R rCG<sup>1</sup> | R LOC | 0.30 | 0.019 | 0.53 | < 0.001 |
| R rCG<sup>1</sup> | R PcG | 0.36 | 0.005 | 0.52 | < 0.001 |
| R rCG<sup>1</sup> | R SFG | 0.35 | < 0.001 | 0.56 | < 0.001 |
| R rCG<sup>1</sup> | R FP | 0.37 | 0.004 | 0.57 | < 0.001 |
| R rCG<sup>2</sup> | L LOC<sup>1</sup> | 0.41 | 0.001 | 0.51 | < 0.001 |
| R rCG<sup>2</sup> | L LOC<sup>2</sup> | 0.49 | < 0.001 | 0.35 | 0.007 |
| R rCG<sup>2</sup> | R LOC | 0.20 | 0.117 | 0.52 | < 0.001 |
| R rCG<sup>2</sup> | R PcG | 0.56 | < 0.001 | 0.58 | < 0.001 |
| R rCG<sup>2</sup> | R SFG | 0.45 | < 0.001 | 0.58 | < 0.001 |
| R rCG<sup>2</sup> | R FP | 0.52 | < 0.001 | 0.62 | < 0.001 |

Pearson's correlation coefficients (r) and corresponding p values describe the correspondence between dog and human ROI dissimilarities for pairs of bodily anger and happiness stimuli. ROIs were identified using the RSA models presented in Tables 4 - 6. R = right; L = left; cESG = caudal ectosylvian gyrus; rESG = rostral ectosylvian gyrus; rSG = rostral sylvian gyrus; mSSG = mid suprasylvian gyrus; cSG = caudal sylvian gyrus; cSSG = caudal suprasylvian gyrus; IC = insular cortex; rCG = rostral composite gyrus; OP = occipital pole; LOC = lateral occipital cortex; PcG = paracingulate gyrus; SFG = superior frontal gyrus; FP = frontal pole. Superscripts 1 and 2 distinguish different ROI peaks within the same anatomical region.

## Discussion

Dogs live immersed in human emotional signals, yet most of what we know about their visual emotion perception comes from their perception of and responses to faces. Here, we provide the first direct neural evidence that the dog brain represents affective information conveyed by human bodies. Across univariate and multivariate analyses, our results converge on complementary findings: an occipito-temporal sensitivity to human bodies in both dogs and humans, emotional modulation in the right caudal suprasylvian gyrus (cSSG) of dogs and the right extrastriate body area (EBA) of humans, and multivariate response patterns that differentiate bodily happiness from anger in both species. Univariate responses to bodily happiness and anger showed substantial spatial overlap in both species, although this overlap appeared less extensive in humans. Notably, the multivariate response patterns suggest similarities in the representational relationships among the bodily emotions, despite differences in the anatomical distribution of these effects: representation of bodily happiness involved more anterior and distributed regions in dogs than in humans, where they were restricted to the right occipital pole, whereas representations of bodily anger were much more extensive in humans than in dogs, where they were restricted to the right rostral composite gyrus. These results suggest similar representational relationships between bodily emotions in dogs and humans, although not necessarily equivalent representations or regions.

### Similar body sensitivity and emotional modulation in the dog cSSG and human EBA

Our findings extend a growing dog fMRI literature showing that the occipito-temporal cortex supports the processing of socially relevant visual stimuli, including animate agents (Boch et al., 2023), faces (Bunford et al., 2020; Cuaya et al., 2016; Dilks et al., 2015; Karl, Boch, Zamansky, et al., 2020; Thompkins et al., 2018), and social interactions (Karl et al., 2021). Within this broader network, the cSSG has previously been identified as body-sensitive in dogs using images of headless bodies (Boch et al., 2023). Here, we extend this evidence to dynamic, full-body human expressions, with the cSSG in dogs and the EBA in humans showing stronger responses to human bodies than to phase-scrambled controls in both hemispheres. Beyond body sensitivity, the right cSSG response was modulated by emotional content, with stronger response to bodily emotions (happiness and anger) than to bodily neutral, suggesting that this region contributes to processing affective information conveyed by human bodies. A similar emotional modulation was found in the right EBA, where bodily emotions have been shown to modulate its response (Peelen et al., 2007; Ren et al., 2022; Ross et al., 2019; Smekal et al., 2026; Van den Stock et al., 2014). Together, our results show that the perception of human bodies relies on visual systems tuned to socially relevant body information in both dogs and humans, and suggest that the right cSSG in dogs and the right EBA in humans are sensitive not only to the visual form of human bodies but also the affective signals they convey (Figure 2, Tables 2 and 3).

### Distributed sensitivity to bodily emotions

The univariate whole-brain contrast of bodily emotions (happiness and anger) against controls (bodily neutral, phase-scrambled happiness, and phase-scrambled anger) revealed distributed regions with stronger responses to bodily emotions (Table 2). In dogs, the responses were primarily located in the temporal lobe but also extended into parietal, occipital, and insular cortices. In humans, the same contrast showed distributed responses predominantly in occipito-temporal regions, extending into parietal, frontal, and medial temporal cortices. This pattern is consistent with previous evidence that visual processing of socially relevant stimuli including human bodies, involves distributed brain regions in dogs and humans (for reviews see Boch, Huber, et al., 2024; Smekal et al., 2026). Importantly, these contrasts included both bodily neutral and phase-scrambled controls, suggesting that the response was not driven simply by the presence of a moving human body or by low-level visual properties.

Within the broader network sensitive to bodily emotions, multivariate response patterns differentiated bodily happiness from bodily anger (Figure 4, Table 4). In dogs, this differentiation was observed in the right ectosylvian and sylvian gyri, and in the left mid suprasylvian gyrus. In humans, the same model showed differentiation in the right occipital pole and lateral occipital cortex. This valence discrimination was absent for the phase-scrambled controls in both dogs and humans, supporting the interpretation that the effect is unlikely to reflect differences in low-level visual properties. These multivariate differentiation of bodily happiness and anger is consistent with previous human evidence that body-sensitive regions can carry distinct multivariate patterns related to different bodily emotions (Ren et al., 2022; Yang et al., 2018). Together, the univariate and multivariate results suggest that the brains of dogs and humans are sensitive to the presence of emotional information in human bodies and also to distinctions between bodily happiness and bodily anger.

### Overlapping responses and asymmetric representations of bodily happiness and anger

Although bodily happiness and bodily anger could be differentiated at the multivariate level, their univariate responses showed substantial overlap in dogs. Separate GLM contrasts for bodily happiness and bodily anger against their respective controls recruited regions primarily located along the temporal lobe, including the suprasylvian, ectosylvian, and sylvian gyri (Figure 3, Table 2). Happiness elicited broader and more anterior response, and anger relatively more posterior response, the BOLD signal extracted from the peaks of both contrasts indicated similar response magnitudes across emotions (Figures S1 and S2). In humans, the same GLM contrasts also showed overlapping responses, particularly in bilateral occipito-temporal regions, although this overlap was less extensive than in dogs. Bodily happiness showed a broader dorsal and sensorimotor distribution, whereas bodily anger showed additional responses in medial and anterior regions (Figure 3, Table 2). These occipito-temporal responses are consistent with previous human fMRI evidence of emotional modulation in body-sensitive visual regions, during dynamic bodily happiness and anger (Peelen et al., 2007; Ross et al., 2019). Parietal and sensorimotor responses to happiness are consistent with human fMRI finding linking bodily happiness to parietal and motor-related regions (Ren et al., 2024), whereas anterior temporal and medial frontal responses to anger align with findings related to affective evaluation and action preparation (Pichon et al., 2008, 2009). This partial overlap suggests that emotion-specific information may not be fully represented by differences in activation magnitude, but may instead be encoded in finer-grained multivariate patterns across overlapping regions.

Within the bodily emotions mask, RSA tested for high within-category similarity and distinct response patterns relative to the other emotion and neutral bodies, revealing significant effects for happiness but not anger in both dogs and humans (Figure 5, Table 5). In dogs, patterns distinguishing happiness from anger and neutral bodies involved the right insular cortex, right caudal sylvian gyrus, right rostral ectosylvian gyrus, and suprasylvian gyri. The involvement of the insular cortex is consistent with previous dog fMRI studies of social and emotional processing (Karl, Boch, Zamansky, et al., 2020; Karl et al., 2021). The sylvian gyrus finding also aligns with evidence that happy human faces elicit distinct multivariate patterns relative to negative expressions such as anger (Hernández-Pérez et al., 2026), suggesting partially shared neural substrates for visual positive affect across facial and bodily channels, consistent with diffusion MRI evidence linking the sylvian cortex to sensorimotor integration of socially relevant communicative cues (Cordeau et al., 2025). In humans, happiness-related patterns were restricted to the right occipital pole, consistent with previous human fMRI studies showing occipital responses to happy bodily expressions (Peelen et al., 2010; Ross et al., 2019). In contrast, the corresponding analysis for anger and the phase-scrambled controls RSA models showed no significant results in dogs or humans. The emotion-specific RSA results therefore suggest that bodily happiness elicits consistent and distinct neural patterns in both dogs and humans, involving more anterior and distributed regions in dogs. The absence of comparable results for bodily anger suggests that its neural representation may follow a different pattern, not captured by the within-condition similarity required by this analysis.

To examine whether anger might follow a different representational pattern, we used whole-brain RSA without requiring within-condition similarity for anger, testing whether it differed from happiness and neutral, which were expected to be similar (Figure 6, Table 6). In dogs, this analysis identified a significant cluster in the right rostral composite gyrus (rCG), this region has been only sparsely characterized in task-based dog fMRI. Resting-state research places the right rCG within a right-lateralized network including secondary somatosensory cortices, the caudate nucleus, and the rostral suprasylvian gyrus (Szabó et al., 2019), while asymmetry findings associate the composite with social action and interaction (Barton et al., 2023). Although the rCG has not been identified as a core in the body-or action-responsive networks in dogs (Boch et al., 2023; Boch, Karl, et al., 2024), it has been implicated in responses to socially relevant human stimuli and learned auditory-reward cues (Cuaya et al., 2023; Karl, Boch, Zamansky, et al., 2020). Thus, this effect may reflect associative, somatosensory, or integrative representation of bodily anger rather than a stable category-specific response pattern within the occipito-temporal body-emotion network. In humans, the same analysis identified a much more extensive pattern involving bilateral lateral occipital cortex, additional occipital, parietal, sensorimotor, and insular regions, as well as the right paracingulate gyrus, superior frontal gyrus, and frontal pole. This distribution is consistent with studies showing that dynamic bodily anger engages affective and action-related systems, including premotor and motor-preparation circuits, possibly reflecting an action-oriented response to interpersonal threat (Kret et al., 2011; Pichon et al., 2008, 2009). In contrast, the corresponding analyses for happiness and the phase-scrambled controls showed no significant results in either species. Overall, the neural differentiation of bodily anger appears to extend beyond established visual body-processing networks in both dogs and humans.

Taken together, the complementary RSA models suggest a similar asymmetry in the representational relationships between bodily happiness and anger in dogs and humans, despite differences in their anatomical distribution. In both species, happiness showed consistent neural patterns distinct from anger and neutral, whereas anger was differentiated from these conditions without requiring within-condition similarity. However, the anatomical distribution differed: happiness involved more anterior and distributed regions in dogs, but was restricted to the right occipital pole in humans, whereas anger involved a much broader network in humans than in dogs. In dogs, this pattern raises the possibility that bodily happiness is particularly relevant for visual social and emotional processing, while anger may involve more contextual or integrative representations. In humans, the broader representation of anger may reflect the involvement of distributed visual, affective, and motor systems.

Unlike some previous dog fMRI studies on human facial emotions, we did not observe significant caudate nucleus involvement during the processing of bodily happiness in dogs. The caudate nucleus has been repeatedly implicated in reward processing and responses to socially relevant cues in dogs, including food-reward prediction, reward representation, familiar human scents, and human happy or emotionally salient faces (Berns et al., 2013, 2015; Cook et al., 2014; Cuaya et al., 2023; Hernández-Pérez et al., 2026; Karl, Boch, Zamansky, et al., 2020; Thompkins et al., 2021). Together with behavioral evidence that dogs attend more to human emotional bodies than to human emotional faces, whereas humans show the opposite attentional preference (Correia-Caeiro et al., 2021), our results raise the possibility that bodily happiness may engage visual systems involved in social and emotional processing without necessarily recruiting reward-related caudate activity. Alternatively, positive faces of humans may signal pending rewards to dogs and thus engage reward anticipation and wanting associate with caudate reward area more robustly than bodies.

### Bodily emotion processing in dogs and humans: a comparative perspective

Our findings can be situated in a broader comparative context. Using the same experimental design in dogs and humans, and a largely matched MRI scanning setup, we observed distributed neural responses to human bodily happiness and anger, despite differences in their anatomical distribution. This pattern is consistent with previous evidence for a distributed network supporting bodily emotion perception in humans, which includes body-selective visual areas, temporal regions involved in biological motion, parietal and premotor systems linked to action preparation, and affective salience regions including the amygdala and insula (Bachmann et al., 2018; De Gelder, 2006; De Gelder et al., 2015; Peelen et al., 2007; Smekal et al., 2026). It also converges with evidence from macaques, where body-selective patches respond more strongly to emotional than neutral bodies across different stimulus species (Taubert et al., 2022). Together, the converging evidence from dogs, humans, and macaques suggest that distributed sensitivity to bodily affective cues may be a broadly shared feature of mammalian social perception, although the spatial distribution of these responses differs between dogs and humans.

A central finding from our direct comparison is that the representational asymmetry between bodily happiness and anger was present in both dogs and humans. In both species, happiness was differentiated from anger and neutral in an RSA analysis requiring within-condition similarity, whereas anger was differentiated in an analysis without this requirement. However, the anatomical distribution differed: happiness involved more anterior and distributed regions in dogs but was restricted to the right occipital pole in humans, whereas anger involved a much broader network in humans than in dogs. Thus, similar representational relationships between bodily emotions were observed in dogs and humans, despite differences in their anatomical organization, highlighting the importance of considering both neural response patterns and anatomy in comparative emotion research (Adolphs, 2026). These differences raise the possibility that bodily happiness and anger may have different relevance for dogs and humans. In humans and macaques, studies on bodily emotion perception have largely emphasized expressions of threat, such as fear and anger, which engage body-selective visual areas, affective regions, and motor cortices (Grèzes et al., 2007; Kret et al., 2011; Pichon et al., 2008, 2009; Taubert et al., 2022). This emphasis is consistent with the broader comparative focus on defensive and aggressive emotional behaviors (Adolphs, 2026). The more extensive responses associated with anger in humans align with this literature, while the more distributed responses to happiness in dogs may reflect the relevance of positive human emotional signals for affiliative interactions. The neural differentiation of bodily happiness in dogs converges with dog fMRI evidence from human facial expressions, where happy faces elicited distinct multivariate patterns relative to negative emotions (Hernández-Pérez et al., 2026). This finding is particularly interesting in light of evidence that dogs’ responses to human happiness cues may depend on social experience. In an odor-based study, human happiness chemosignals did not differentiate puppies’ behavior from the control condition but did elicit distinct behavioral patterns in adult dogs, the authors suggested that responses to happiness chemosignals may require learning during early socialization (D’Aniello et al., 2023). In our study, the neural differentiation of bodily happiness in dogs may therefore reflect the relevance of human happiness for affiliative interactions in adult companion dogs. Another, not mutually exclusive, possibility is that our participants have more frequent or more consistent experience with humans expressing happiness than anger toward them. Bodily anger, by contrast, may be more context dependent, particularly in a silent visual paradigm with unfamiliar actors and blurred faces, where dogs lacked vocal, facial, olfactory, and familiar-context cues. Angry expressions from unfamiliar actors may also have been less salient than those displayed by their caregivers.

The direct comparison of stimulus-pair dissimilarities further suggests that dogs and humans share aspects of the representational organization of bodily happiness (Figure S3, Table 7). Across most ROI comparisons, the dissimilarities among bodily happiness stimuli corresponded more closely between dogs and humans than those among bodily anger stimuli. The weaker correspondence for bodily anger suggests that its representational organization may differ more between dogs and humans.

## Limitations

Some limitations should be considered when interpreting the study’s results and their relevance for future research. First, although full-body videos increased ecological validity relative to static images, our stimuli were unimodal, portrayed by unfamiliar actors, and presented outside a natural interactional context, which may not fully capture the perception of human emotions in everyday multimodal interactions. Second, while previous behavioral studies show that dogs can discriminate human emotional cues across sensory modalities, our stimuli, as in many studies of emotion perception, were validated by human observers. These human-defined categories may not map perfectly onto dogs’ own perceptual or affective representations. Third, we tested only two bodily emotions, happiness and anger. This choice was based on behavioral evidence that dogs discriminate between these two categories in human facial expressions, but including other emotions such as fear, which is more extensively characterized in the human and primate bodily emotion literature, would provide a more comprehensive picture in future studies. Fourth, our sample consisted of highly trained companion dogs, which may limit generalizability to the broader dog population. Factors such as self-selection, training history, rearing background, acclimation, and experience with humans may influence the generalizability of our findings (Webster & Rutz, 2020), a point that is particularly relevant for interpreting the happiness-related multivariate patterns observed in our results. Fifth, our stimuli were specifically designed as human emotions directed toward dogs, including situations involving dog-human interactions and a camera perspective approximating dogs’ eye level. Although this design allowed us to present the same stimuli to dogs and humans, these expressions may be perceived differently from emotions directed toward other humans. Future studies combining multimodal naturalistic stimuli, direct behavioral measures, and a broader range of emotional categories will be essential to determine how dogs integrate bodily, facial, vocal, olfactory, and contextual cues when perceiving human emotions, and how these processes compare with those in humans. Finally, because we presented only human bodily expressions, our design does not establish whether dogs represent emotions conveyed by conspecific bodies in a similar way. Future comparisons of conspecific and heterospecific bodily emotions would help clarify how stimulus species influences emotional representations in dogs.

## Conclusion

Our study highlights the added value of a shift from the investigation of dogs’ visual emotion perception from faces to an emotional channel that is central in humans and may be especially informative for dogs: the body. Using fMRI data from 24 awake companion dogs and 40 humans viewing the same dynamic human bodily emotional expressions and control stimuli, we provide the first neural evidence of how the dog brain represents affective information conveyed by human bodies. Specifically, both dogs and humans showed occipito-temporal sensitivity to human bodies, emotional modulation in body-sensitive visual regions, along with distributed multivariate response patterns that differentiated bodily happiness from anger. Bodily happiness elicited consistent category-specific response patterns in dogs and humans, involving more anterior and distributed regions in dogs and the right occipital pole in humans. In contrast, bodily anger was distinguishable under a model that did not require high within-condition similarity, involving the right rostral composite gyrus in dogs and a much more extensive occipital, parietal, sensorimotor, insular, and frontal network in humans. Together, these findings suggest that dog and human brains show similar representational asymmetries but different neural distributions for bodily happiness and anger. More broadly, our results advance comparative social neuroscience and highlight bodily expressions as an important channel for investigating emotional communication between dogs and humans.

## Data availability

The data supporting the findings of this study will be made publicly available on Zenodo upon publication of the article.

## Code availability

The code is available on: https://github.com/rhernandez00/dog_brain_toolkit

## Acknowledgements

We thank the dogs and their families for their time and enthusiasm in participating in this project. We are also very grateful to Laura Lausegger and Marion Umek for their dedicated work in training the dogs and supporting data collection, together with Katharina Fiala, Max Rosenthaler, Sara Binder, and Zöe Hauger with the support of Johannes Hoops, Katharina Langer, Liene Kruskopa, Louisa Pillwax, Matthias Jung, Nino Sidar, Radek Sharma, and Tamara Puchegger, at the Comparative Canine Neuroimaging Unit. We further acknowledge Karin Bayer, the actresses and actors, and the survey participants for their valuable assistance in recording and validating the videos used as stimuli in this project.

## Author contributions

**LVC:** Conceptualization, Methodology, Formal analysis, Investigation, Data Curation, Writing - Original Draft, Writing - Review & Editing, Visualization, Project Administration. **RH-P:** Conceptualization, Methodology, Software, Validation, Formal analysis, Investigation, Data Curation, Writing - Review & Editing, Visualization. **MB:** Methodology, Resources, Writing - Review & Editing. **JR:** Methodology, Formal analysis, Writing - Review & Editing, **AR-A:** Methodology, Writing - Review & Editing. **LH:** Conceptualization, Methodology, Resources, Writing - Review & Editing, Supervision, Funding acquisition. **CL:** Conceptualization, Methodology, Resources, Writing - Review & Editing, Supervision, Funding acquisition.

## Competing interest

The authors declare no competing interests.

## Funding

This research was funded in whole or in part by the Austrian Science Fund (FWF): 10.55776/ESP602, 10.55776/J4828, and “DK Cognition and Communication 2”: W1262-B29 [10.55776/W1262].

For open access purposes, the author has applied a CC BY public copyright license to any author accepted manuscript version arising from this submission.

## Notes

### Competing Interest Statement

The authors have declared no competing interest.

